# Increased rates of hybridization in swordtails are associated with water pollution

**DOI:** 10.1101/2025.04.22.649978

**Authors:** Benjamin M Moran, Wilson F Ramírez-Duarte, Daniel L Powell, Terrance TL Yang, Theresa R Gunn, Gaston I Jofre-Rodríguez, Cheyenne Y Payne, Erik NK Iverson, Gabriel A Preising, Shreya M Banerjee, Alexandra E Donny, Rhea Sood, John J Baczenas, Gabriela María Vázquez Adame, Carla Gutiérrez-Rodríguez, Chelsea M Rochman, Molly Schumer, Gil G Rosenthal

**Affiliations:** Department of Biology, Stanford University, 327 Campus Drive, Stanford, CA 94305, USA; Centro de Investigaciones Científicas de las Huastecas “Aguazarca”, A.C., 16 de Septiembre, 392 Barrio Aguazarca, Calnali, Hidalgo 43240, México; Department of Evolution and Ecology, University of California at Davis, University of California, Davis, 475 Storer Mall, Davis, CA 95616, USA; Department of Ecology & Evolutionary Biology, University of Toronto, 25 Willcocks Street, Room 3055, Toronto, Ontario, M5S 3B2, Canada; Present Address: Department of Environmental & Physical Sciences, Concordia University of Edmonton, 7128 Ada Blvd NW, Edmonton, AB T5B 4E4, Canada; Department of Biological Sciences, Louisiana State University, 386 S Campus Dr., Baton Rouge, LA 70802, USA; Department of Biology, Virginia Commonwealth University, 1000 W Cary St, Richmond, VA 23284, USA; Department of Ecology and Evolutionary Biology, University of California, Santa Cruz, 130 McAllister Way, Santa Cruz, CA 95060, USA; Southwest Fisheries Science Center, National Marine Fisheries Service, National Oceanic and Atmospheric Administration, 110 McAllister Way, Santa Cruz, CA 95060, USA; Department of Integrative Biology, University of Texas at Austin, 2415 Speedway, Austin, TX 78712, USA; Department of Genome Sciences, University of Washington, 3720 15th Ave NE, Seattle WA 98195, USA; Red de Biología Evolutiva, Instituto de Ecología A.C., Carretera antigua a Coatepec 351, Col. El Haya, Xalapa, Veracruz 91073, México; Freeman Hrabowski Fellow, Howard Hughes Medical Institute, 4000 Jones Bridge Road, Chevy Chase, MD 20815, USA; Dipartimento di Biologia, Università degli Studi di Padova, Via Ugo Bassi, 58/B, Padova, 35131, Italy

**Keywords:** hybridization, hybrid populations, anthropogenic change, water quality, mate choice, freshwater fish, Poeciliidae, land use

## Abstract

Understanding the nature of reproductive barriers separating species is a fundamental goal of evolutionary biology. Such barriers may be environmentally sensitive, and recent research has documented an increasing number of cases where anthropogenic environmental disturbance is associated with hybridization. However, few studies have been able to quantify and compare potential environmental mechanisms connecting anthropogenic disturbance to hybridization. Here, we combine genomic and environmental surveys to explore the loss of reproductive isolation between *Xiphophorus malinche* and *X. birchmanni*, fishes whose riverine habitat in montane Mexico is increasingly impacted by human-mediated disturbance. By inferring genome-wide ancestry in thousands of fish, we characterize the landscape of hybridization between these sister species in four drainages. Ancestry structure varies across streams from stable coexistence to clinal hybrid zones, hinting that hybridization dynamics in this system may be environmentally dependent. In one stream, sites upstream of an urbanized area host distinct sympatric ancestry clusters, while downstream sites collapse into a hybrid swarm. By sequencing mothers and embryos, we show that assortative mating is weakened downstream of this urbanized area. We hypothesize that the downstream hybrid swarm is driven by chemical disruption of olfaction that impacts mating preferences. Water chemistry measurements show significant changes across the urbanized area, including in parameters known to disrupt fish olfaction and mating. We identify alterations of the olfactory epithelium between sites upstream and downstream of the urbanized area consistent with differential effects of water quality. Taken together, our work illuminates potential mechanisms linking anthropogenic disturbance to the breakdown of reproductive isolation.

## Introduction

To understand speciation, evolutionary biologists have long studied cases in which reproductive barriers between species are incomplete and hybridization can occur^1,2^. Such studies have sometimes found evidence that hybridization occurs in areas of environmental disturbance. Disturbance can bring together species that do not otherwise coexist, create novel environments where hybrids thrive, or alter the interactions between species that previously coexisted without interbreeding^3–5^. Empirical work has shown that premating barriers, such as a behavioral preference for mating with conspecifics (i.e. assortative mating), are highly context-dependent^6–10^ and sensitive to environmental conditions^11,12^. They may therefore disappear in the face of disturbances that impact mechanisms of mate discrimination^13–15^.

While natural changes to the environment may have been the major source of disturbance historically, human impacts (e.g. land use change, chemical pollution, resource extraction) may now be playing an analogous role^14,16–18^. Human modification of the environment can interfere with the mechanisms by which conspecifics identify each other, either by directly affecting signal production and transmission or by abolishing conspecific preferences^14,19–21^. In aquatic species, in particular, a variety of chemical pollutants have been shown to weaken conspecific mate preferences^14,19,20,22–24^. As human impacts on the environment become ubiquitous^25^, the reported rate of hybridization appears to be increasing across a range of taxa^5^.

If assortative mating is a major reproductive barrier between species, weakening this barrier can lead to a collapse of genetic and phenotypic differentiation^26–30^, often referred to as a “hybrid swarm”. Since mate discrimination is typically weaker in hybrids than in parental species^13^, this can lead to irreversible phenotypic and genetic homogenization through hybridization (i.e. “speciation reversal” or “genetic swamping”). Although hybridization may sometimes lead to adaptive introgression^31,32^ or the formation of new species^33^, the immediate consequence of the formation of hybrid swarms is more often the loss of genetic differentiation and functional biodiversity^18,34^. Moreover, early-generation hybrids often experience intrinsic hybrid incompatibilities^35,36^ and thus hybridization may also decrease fitness and overall population size.

Swordtails (Teleostei: genus *Xiphophorus*) provide a compelling system to study the breakdown of reproductive isolation following anthropogenic disturbance. As model systems for hybridization and mate choice, these stream fishes have been the subject of numerous studies showing that mating preferences can be altered by environmental and social context^7,8,13,19^. Swordtails are found along the eastern slopes of the Sierra Madre Oriental and Gulf Coastal Plain of Mexico, a region whose water quality and biodiversity^37,38^ have been impacted by growing human population density and agricultural land conversion since the 20^th^ century^39,40^. The sister species *X. malinche* and *X. birchmanni* overlap in the montane streams of northern Hidalgo state, Mexico, where steep slopes create frequent migration barriers (waterfalls, cascades, and subterranean stream reaches) and coincide with dramatic ecological variation over short geographic distances. *X. malinche* occupy cool headwaters, while *X. birchmanni* are found in warmer lower-elevation streams, creating the potential for ecological selection against hybrids^41,42^. Genome-wide divergence between the species is moderate (D_xy_ = 0.45%)^43^ and their early-generation hybrids suffer from severe genetic incompatibilities^44–46^. However, some hybrids are viable and fertile, and hybrid populations can be found in multiple independent stream reaches^43,47^. Researchers first reported hybridization between *X. malinche* and *X. birchmanni* in 1997, whereas hybrids were not detected when sampling the same stream reaches a decade earlier^48,49^. Meanwhile, the period of 1980–1995 saw a 37% increase in human population in the state of Hidalgo^50^. Genetic approaches have estimated that these hybrid zones are on the order of ∼100 generations old^51^, consistent with hybridization beginning in the late 1900s. This recent onset of hybridization raises the question of what triggered it, and whether it was tied to the concurrent socio-ecological transformation of the region.

Though we lack data from these streams prior to the start of hybridization, we can still study the environmental factors which affect hybridization rates by observing spatial patterns in present-day populations. The genetic structure in most hybrid populations between *X. malinche* and *X. birchmanni* is best described as a hybrid swarm lacking population structure^43^. One exception is a population just upstream of the town of Calnali, Hidalgo. There, genetic studies have described two subpopulations composed of *malinche*-like and *birchmanni*-like admixed fish that exist in sympatry and co-occur at small spatial scales (Figure 1)^52^. The long-term history of hybridization between these populations is uncertain, but their observed coexistence and partial admixture is consistent with some level of past gene flow counteracted by selection against hybrids^53^. Sequencing over multiple years showed that this bimodal population structure was stable as early as 2003 and is driven by strong assortative mating between individuals of the same ancestry group^52^. However, recent sampling showed that population structure disappears just downstream of the town of Calnali^44^, raising the possibility that assortative mating breaks down after exposure to a point pollution source.

**Figure 1.**
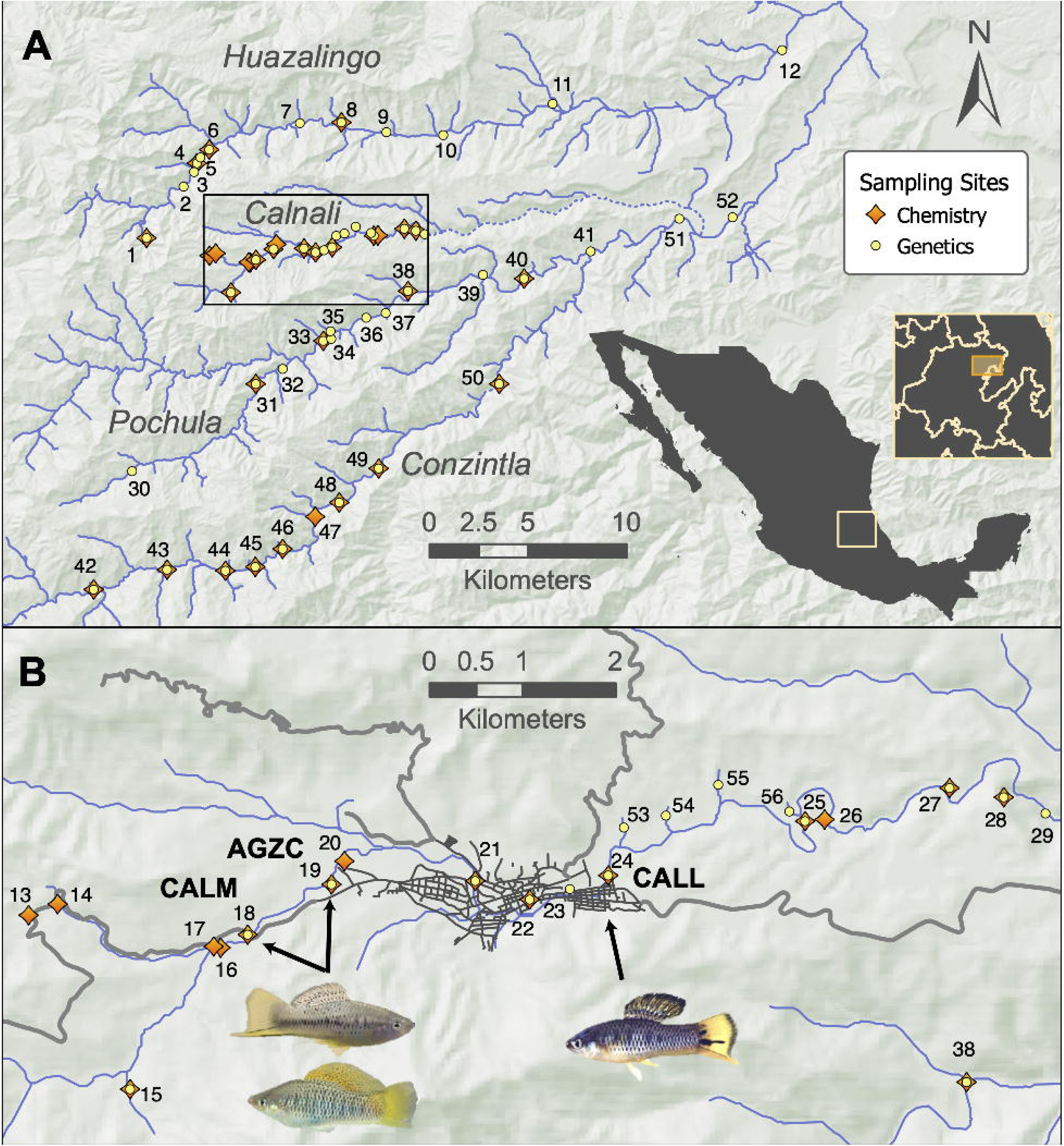
Study area and sampling sites. **A)** Overview of Río Atlapexco basin, highlighted in orange within the inset Mexican state map, whose area is indicated in yellow on the national silhouette. Streams are shown as blue lines, seasonally dry subterranean reaches are dashed, the four major streams studied are labelled, fish collection sites are yellow circles, and water chemistry measurement sites are orange diamonds. The black box encloses the area displayed in panel **B**. **B)** Sites sampled on the Río Calnali. Gray lines show paved roads, and black lines show streets in the town of Calnali. Images show representative male *Xiphophorus* fishes with *X. malinche*-like (top left), *X. birchmanni*-like (bottom left) and admixed (right) phenotypes as observed in three populations on the Río Calnali (AGZC = Aguazarca, CALM = Calnali Mid, CALL = Calnali Low). See also Table S1.

Here, we synthesize biological, geographic, and water chemistry datasets to assess the relationship between environmental conditions and patterns of hybridization across multiple *X. malinche × X. birchmanni* hybrid zones in Hidalgo, Mexico (Figure 2). We use low-coverage whole-genome sequencing of 2620 fish to survey hybrid ancestry in four demographically independent streams and confirm the collapse of population structure around relatively urbanized areas within the Río Calnali, a pattern which is not replicated in other drainages. We then use mother-embryo sequencing in the Río Calnali to confirm that assortative mating by ancestry is weakened downstream of the urbanized area. Next, we document environmental changes which spatially coincide with the unique loss of reproductive isolation within the Río Calnali. Using land use classification of satellite imagery, we confirm that changes in population structure downstream on the Río Calnali are coupled with a high density of built infrastructure near the stream. We also find that population structure in the Río Calnali breaks down at the point where we identify unique shifts in the stream’s physical chemistry, organic and inorganic nutrient loading, and metal concentrations. We then test for similar shifts in metal concentrations in fish tissue. Finally, given the known importance of olfactory communication in swordtail mate choice, we assess olfactory epithelial damage as a potential mechanism linking environmental change to hybridization. We test for variation in olfactory rosette morphology along the Río Calnali, as well as evidence of histological changes after laboratory exposure to humic acid, copper, and ammonia. While we observe no effect of short-term laboratory chemical exposure on olfactory histology, we find evidence of greater histological damage in samples from polluted downstream sites compared to less disturbed upstream sites. This work thus explores potential links between environment, phenotype, and genomics in an anthropogenic hybrid zone, deepening our understanding of human impacts on biodiversity.

**Figure 2.**
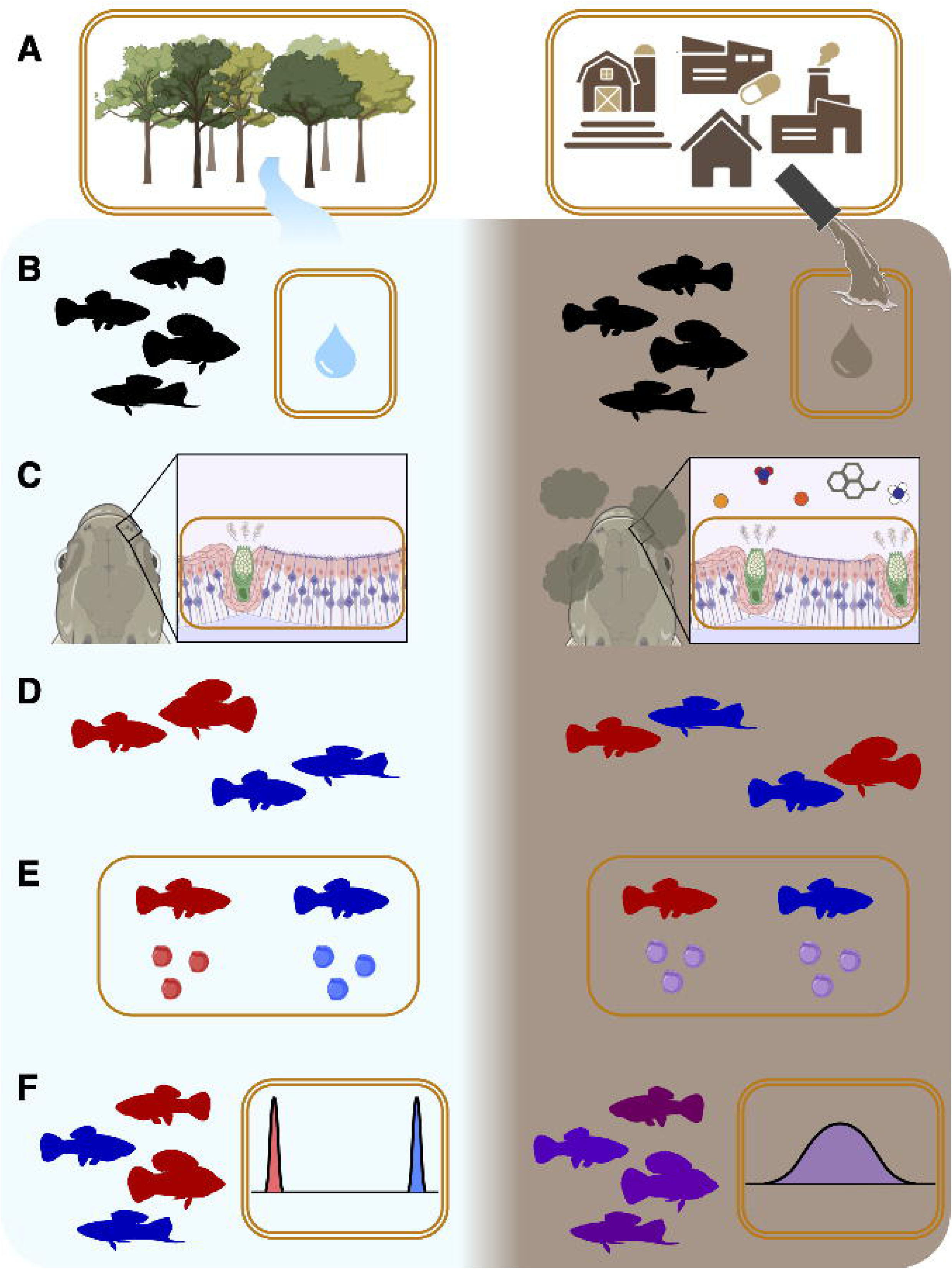
Overview of hypothesized mechanisms connecting human development to swordtail hybridization. Orange boxes represent aspects observed in this study, with double boxes measured across all streams, and single boxes measured only within the Río Calnali. **A)** Levels of anthropogenic development differ across space in the Río Atlapexco drainage, leading to differential runoff which creates **B)** novel variation in chemical environment within and between rivers. **C)** Environmental change affects sensory biology and communication, **D)** attenuating preferences for mates of similar ancestry, and leading to **E)** decreased assortative mating by ancestry and **F)** the collapse of subpopulations into hybrid swarms. Created in BioRender. Schumer, M. (2026) https://BioRender.com/zgh0wna.

## Results

### Population structure across streams

Ancestry structure varied markedly among the four streams sampled in this study. One stream, the Pochula, showed a relatively smooth cline in ancestry from pure *X. malinche* at higher elevations to *X. birchmanni-*like hybrids at lower elevations. The other three streams varied in the degree of population structure, with sites in the Calnali and Conzintla showing significant departures from unimodality in the distribution of individual ancestry proportions (Figure 3; Hartigan’s *D* = 0.12–0.19, *P < 0.0001*). Examination of the variation in individual ancestry in these populations supported the presence of two subpopulations with distinct genome-wide ancestry.

**Figure 3.**
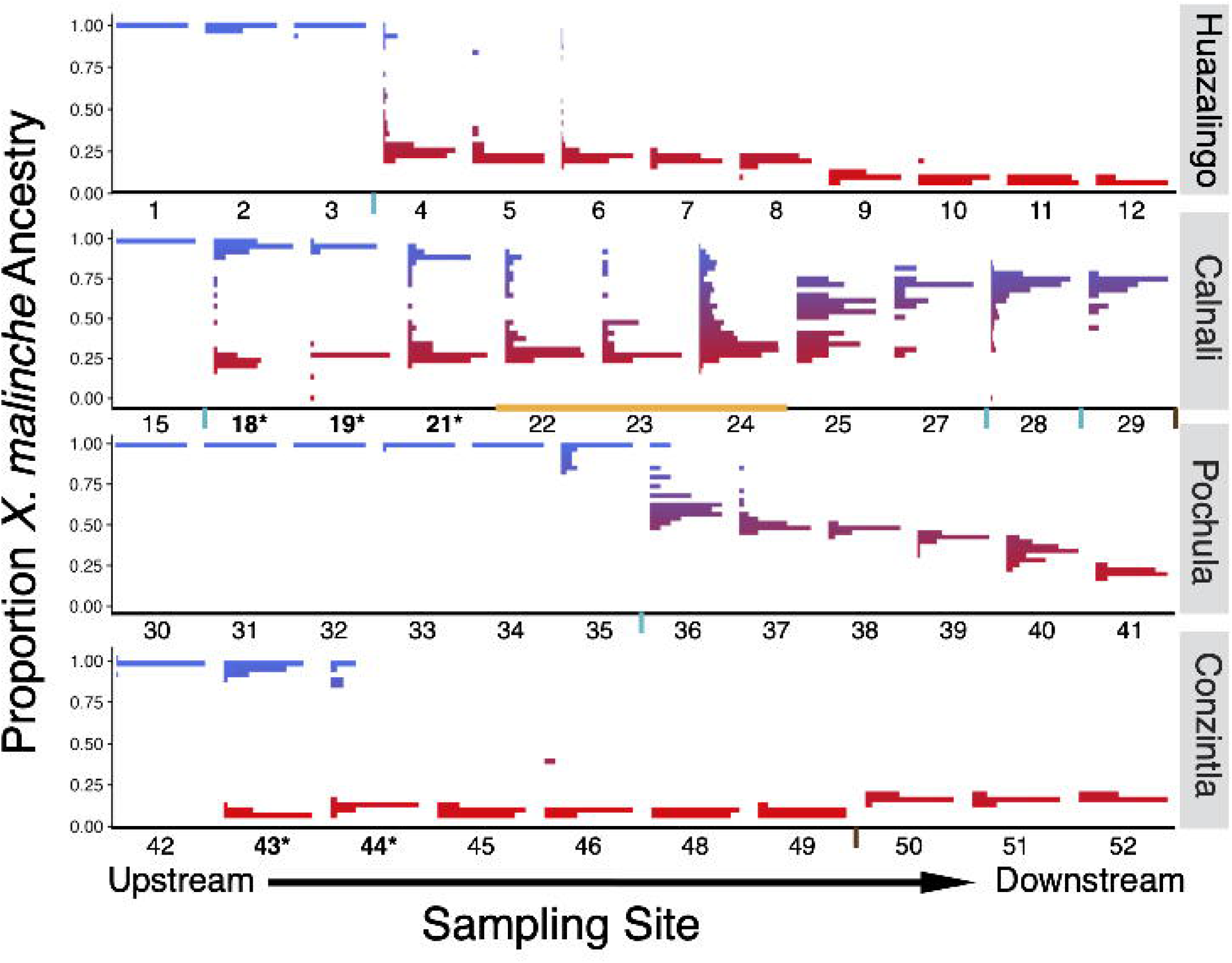
Ancestry distributions of *X. malinche × X. birchmanni* hybrids along four streams. Sites are arranged along the X-axis with upstream sites on the left and downstream sites on the right. Y-axis corresponds to the proportion of genome-wide ancestry derived from *X. malinche* (1 = pure *X. malinche*, 0 = pure *X. birchmanni*), and the width of bars indicates the count of individuals in that ancestry bin in that population (sample sizes per population in Table S3). X-axis text shows site numbers as in Figure 1, with bolding and asterisks indicating sites with significant deviations from a unimodal ancestry distribution (Table S3). Orange segment on the Río Calnali transect denotes the sites located within the town of Calnali. X-axis ticks denote likely migration barriers between sites (blue = dams, waterfalls, and rapids; brown = subterranean stream reaches). See also Figures S1, S8 and Tables S2, S3–S4.

Though multiple streams showed sites with bimodal population structure, we found that the extent of this structure differed starkly across drainages. In the Río Conzintla, we observed sympatry between nearly-pure *X. malinche* and *X. birchmanni* at two sites separated by 3.7 stream kilometers (mean ± standard deviation minor parent ancestry 5.6 ± 3.1% at Xochicoatlán, 10.6 ± 4.5% at Mixtla), with no recent-generation hybrids detected. Downstream of these sites, the *X. malinche* cluster disappeared and left a single *X. birchmanni* ancestry cluster with a spatially stable ancestry fraction (Kolmogorov-Smirnov Test *P* > 0.05; Figure 3; Table S3). Combined with our observation of only 1 recent hybrid out of 239 individuals, this suggested that gene flow between ancestry groups in this stream is limited. In the Río Huazalingo, the presence of two subpopulations was only observed at sites directly downstream of a waterfall separating pure *X. malinche* from admixed *X. birchmanni*, and ancestry structure was not significant after Bonferroni correction (Table S3). *X. malinche-*like individuals were detected up to 0.91 km downstream of the waterfall, but this pattern varied between sampling seasons, hinting that the *X. malinche* individuals may not persist there (Figure S1A).

By contrast, in the Río Calnali, we observed a dramatic collapse in population structure centered on the town of Calnali (Figure 3). Three sites upstream of the town, spanning 3.15 stream kilometers, had significantly multimodal ancestry distributions (Hartigan’s *D* = 0.12–0.21, *P* < 0.00001; Table S3), with *X. malinche*-like and *X. birchmanni*-like subpopulations co-existing in sympatry. Sites downstream of the town plaza showed evidence of a unimodal distribution (*D* = 0.01–0.07, *P* > 0.18 across all sites; Table S3), with intermediate ancestry and increased variance in individual ancestry proportion. Intriguingly, average *X. malinche* ancestry increased downstream of the town (Figure 3), in contrast to expectations given the elevational distribution of *X. malinche* populations in other streams. Sequencing of four tributaries connected to this reach of the Calnali showed that each hosted a single ancestry cluster, with three composed of *X. malinche-*like fish (Figure S1D). The shift towards *X. malinche* ancestry downstream of these tributaries may therefore be a result of propagule pressure.

### Strength of ancestry-assortative mating in the Calnali Low and Aguazarca hybrid populations

In the upstream Calnali Mid site on the Río Calnali, we previously documented bimodal ancestry structure and near-complete assortative mating by ancestry^52^. By contrast, the Calnali Low site does not have significantly bimodal ancestry structure, although variance in ancestry is high (Figure 3). We used paired mother-embryo sequencing to evaluate whether mating patterns differed from expectations under random mating with respect to ancestry. Despite the intervening breakdown in population structure, we found that mothers and offspring in both upstream and downstream populations were less different in ancestry than expected under simulated random mating (*P <* 0.001 by simulation; Figure 4A–B). However, the deviation from neutral expectations was greater at Calnali Mid than at Calnali Low (mean 0.0655 vs 0.0257, *t*-test *P* = 0.0483). This suggested that assortative mating by ancestry is weaker at the Calnali Low site.

**Figure 4.**
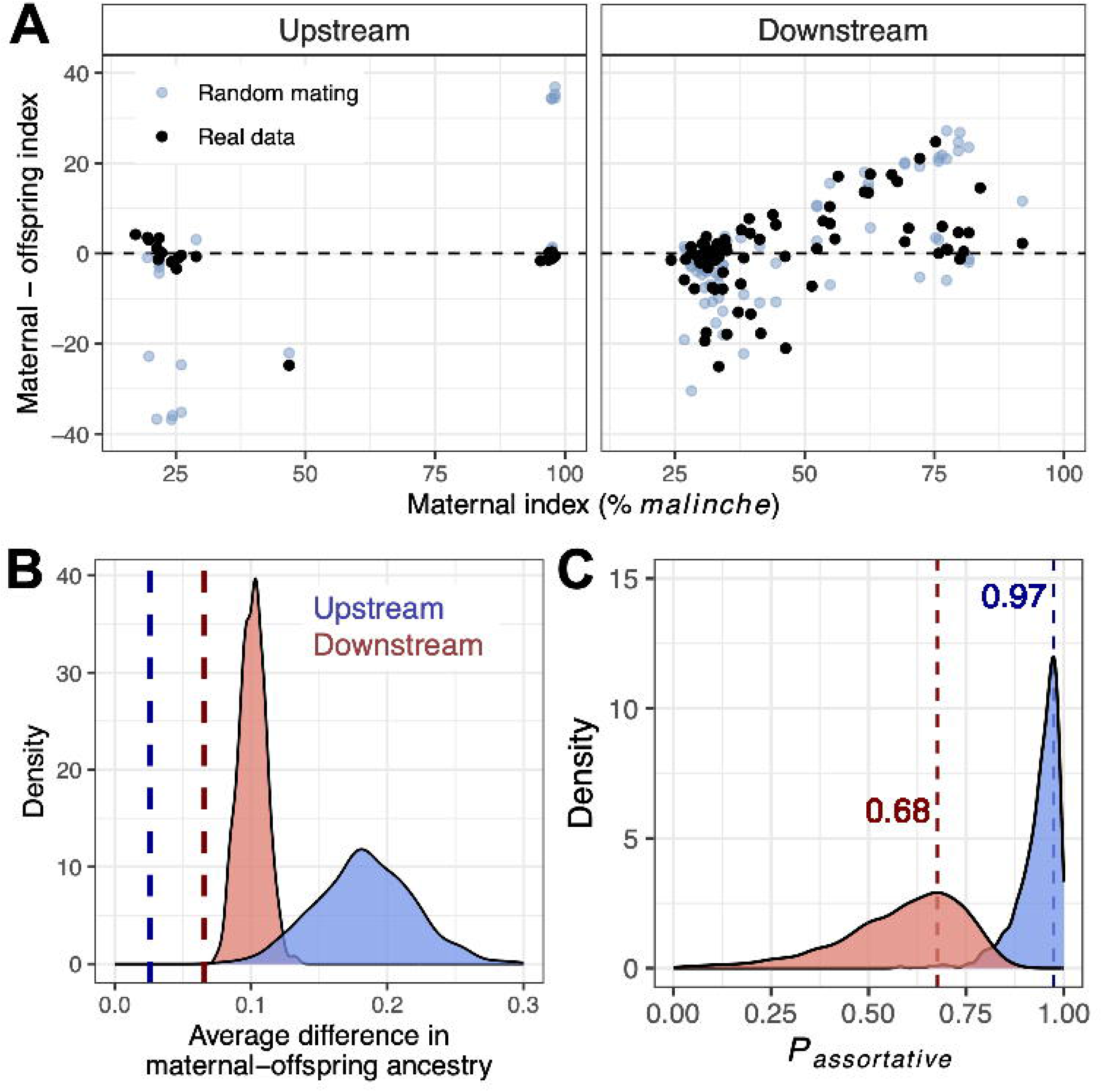
Mating patterns in fish upstream and downstream of the town of Calnali. **A)** Mother-embryo ancestry from upstream (Calnali Mid) and downstream (Calnali Low) of the town of Calnali. X-axis shows the proportion of each mother’s genome derived from *X. malinche*, and Y-axis shows the difference in genome-wide ancestry between each mother and embryo. Black points are observed values, blue point are a simulated sample set under random mating. **B)** Observed mother-embryo ancestry differences diverge from random mating expectations at both sites. Color denotes sampling site, density curves show the distribution of average mother-embryo ancestry difference across 1000 simulations of random mating, and dashed lines show the observed average mother-embryo ancestry difference. Mean ancestry difference was significantly lower than expected under random mating for both sites, consistent with assortative mating by ancestry (*P < 0.001*). **C)** Inferred strength of assortative mating from observed maternal-offspring ancestry distributions. Density curves depict the simulated probability of assortative mating (*P_assortative_*) in accepted simulations for each population, with 0 representing random mating. Dashed line denotes MAP estimates (Calnali Low = 0.68, Calnali Mid = 0.97).

As such, we sought to quantify the strength of ancestry-assortative mating in the Calnali Low population and compare it to the upstream Calnali Mid population. We used an approximate Bayesian computation approach to ask what strength of ancestry-assortative mating was consistent with the average mother-offspring ancestry difference at each site (see Methods). Based on accepted simulations, we recovered a well-resolved posterior distribution of the strength of ancestry-assortative mating (*P*_assortative_), and a maximum *a posteriori* (MAP) estimate for Calnali Low of 0.68 (95% CI 0.20–0.82; Figure 4C), substantially weaker than the estimate from Calnali Mid (MAP = 0.97, 95% CI 0.79–0.99). A direct comparison of the accepted simulations confirmed that assortative mating was significantly stronger at Calnali Mid (Wilcoxon signed rank test *P* < 2.2e-16), with a MAP estimate for the difference in the strength of assortative mating of 0.27 (95% CI 0.08–0.72). Thus, our results suggested that weaker assortative mating contributes to the breakdown in population structure in the Calnali Low hybrid population.

### Land use across drainages

Land use quantification showed that human impacts on the Río Calnali watershed were higher than other streams. The reaches where the stream passed through the town of Calnali had the highest proportion of developed land in the study area, with increased levels of urbanization centered on the location of population structure collapse (Figure 5). When calculating land use in the cumulative watershed of each site, the downstream sites on the Río Calnali had the highest proportions of developed land in the study area (Figure 5C), especially when considering land within a short distance of the stream (Figure 5E). Associations between cleared land (vegetation, with the exclusion of forest) and population structure within the Río Calnali were less clear (Figure S2), but the downstream section of the Río Calnali did show the highest cumulative cleared area close to the stream among all reaches surveyed (Figure S2F, S2H). Overall, our findings suggested that some degree of deforestation and development is ubiquitous throughout the Atlapexco drainage, but the level of built infrastructure was especially high in the Calnali subcatchment.

**Figure 5.**
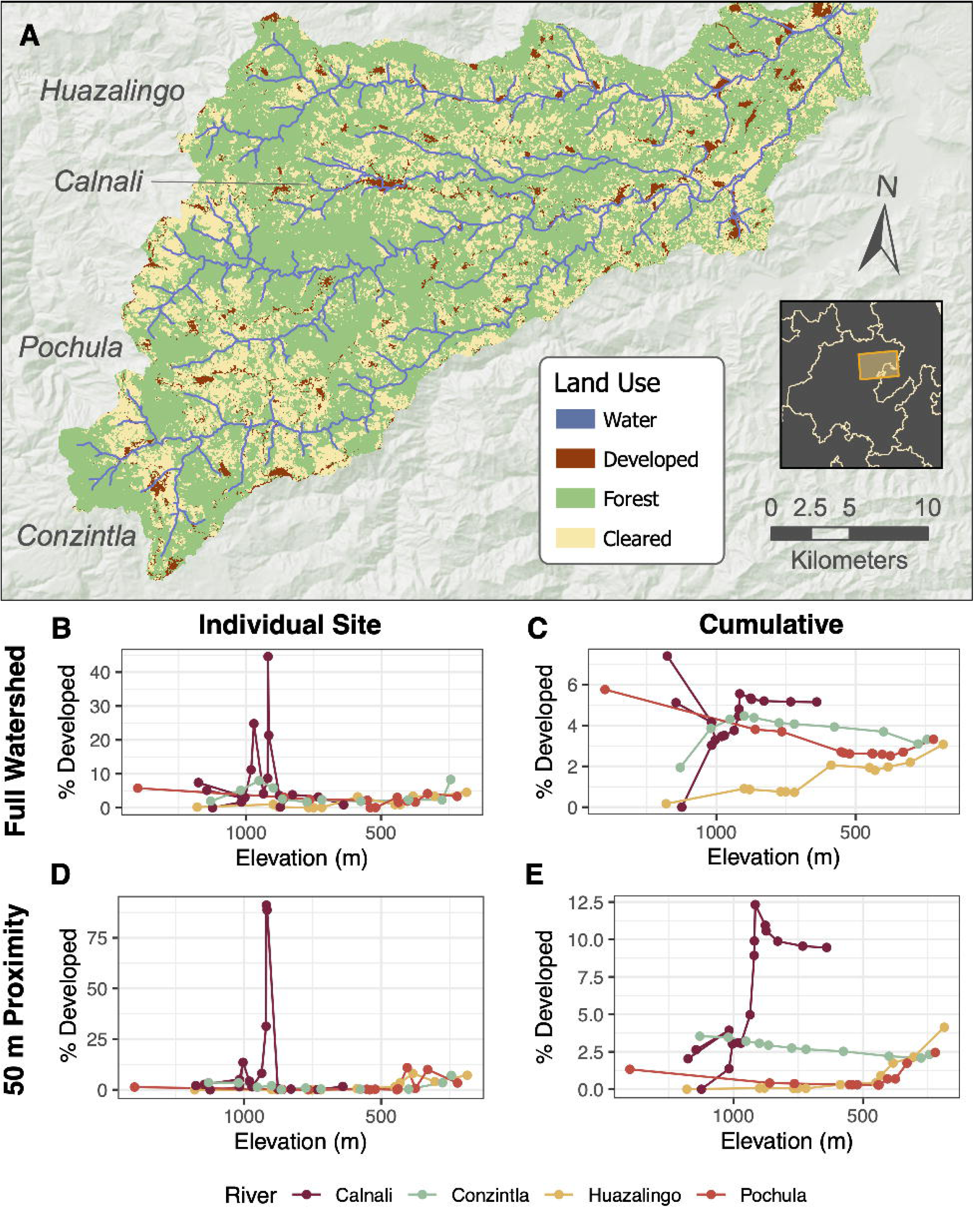
Land use classification analysis in the Río Atlapexco basin. **A**) Map of classifications from 2020–2023 hyperspectral imagery. 10 m^2^ pixels were classified as developed (buildings, mines, and roads), cleared land (grassland, pasture, or cultivated crops) forest (continuous tree cover, excluding plantations), or water by an SVM algorithm, with training polygons identified from 3 m^2^-resolution 2022 imagery from Planet Labs^101^. Stream polygons are shown as blue lines over classified pixels. **B–E)** Percent of land classified as developed within different drainages of the Río Atlapexco basin. Points show the proportion of developed pixels in each site’s subcatchment. Left column (**B** and **D**) uses the subcatchment of the section of stream between a site and the closest upstream site. Right column (**C** and **E**) uses the cumulative subcatchment of all sections of the stream upstream of a given site. Top row (**B** and **C**) shows calculations that include pixels of any distance from a stream, while bottom row (**D** and **E**) shows calculations limited to pixels within 50 m of a stream. Color denotes drainage, and lines connect sites in the same stream based on their order. The line corresponding to the Río Calnali branches because multiple headwater tributaries were sampled. See also Figure S2.

### Variation in water and tissue chemistry

Water chemistry varied starkly both between streams and between sites on the Río Calnali. Principal component analysis demonstrated that most variation in water quality could be summarized by two axes: one that largely separated the sites by stream (PC1), and another separating the sites upstream of the town of Calnali from the other streams and those downstream of Calnali (PC2; Figure 6). Conductivity, alkalinity, hardness, and pH varied across streams, with sites in the Río Conzintla having the highest values and sites in the Río Calnali having the lowest (Figures 6A, S3). Sites downstream of the town of Calnali were associated with increased dissolved organic matter, turbidity, ammonia, and nitrite, as well as reduced dissolved oxygen, all consistent with increased anthropogenic impacts (Figure 6A). Statistical tests identified significantly elevated levels of turbidity, DOM, nitrite, and ammonia in the sites downstream on the Río Calnali relative to the upstream sites and to other streams (Figure 6C–D, Table S5).

**Figure 6.**
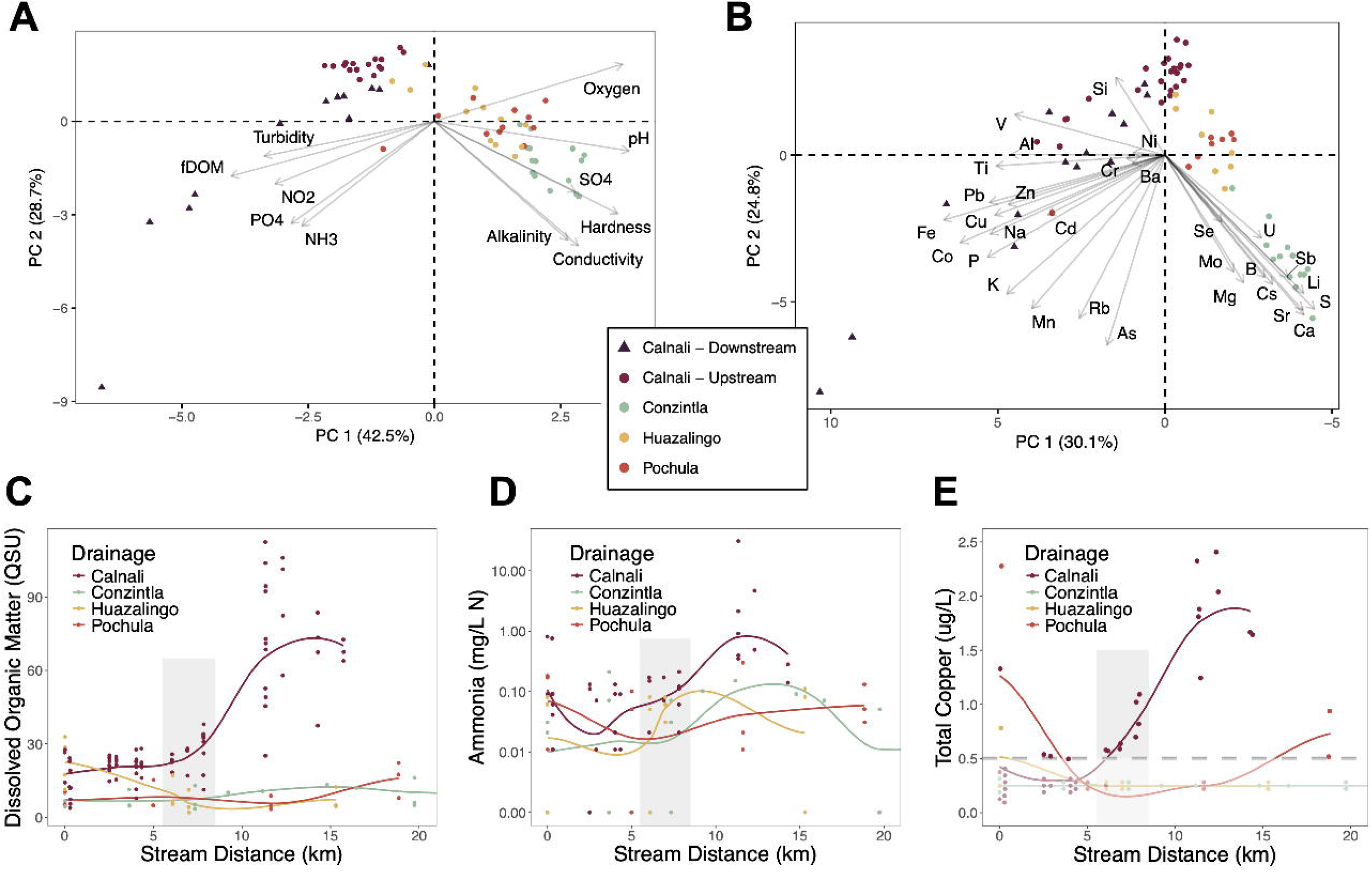
Water chemistry data from four streams in the Río Atlapexco drainage. **A–B)** Principal component analysis (PCA) biplots for **A)** parameters measured using the sonde, turbidimeter, colorimeter, and **B)** parameters measured using ICP-MS. Points represent sampling events, and color and shape denote drainage, with sites on the Río Calnali upstream and downstream of the town of Calnali labelled separately. Labelled arrows represent variable loadings onto the first two principal components. **C–E)** Observations of water chemistry parameters elevated in sites downstream of the town of Calnali, including **C)** fluorescent dissolved organic matter, **D)** ammonia, and **E)** total copper. Grey shaded area denotes the sites located within the town of Calnali on the Río Calnali transect. X-axis indicates distance downstream from the first site sampled on a stream, points indicate individual measurements, and lines show a LOESS-spline fit of change over the stream course. Note that the Y-axis for ammonia (**D**) is log-scaled. Translucent points below the gray dashed line in **E)** represent measured values below the detection limit of the ICP-MS assay, which are plotted at arbitrary Y-values and only represent sample counts. See also Figures S3–S6, Table S5, and Data S1.

Turning to ICP-MS measurements of metal ion concentrations, a principal component analysis showed that concentrations of many ions were highest in the Conzintla and lowest in the Calnali (Figures 6B, S3), potentially due to hydrological and geological variation (Figure S4)^54^. The transition from the sites upstream to downstream of the town of Calnali was also associated with a dramatic increase in the concentration of numerous ions, especially major industrial metals (e.g. iron, aluminum, copper, lead, manganese; Figure 6B). Multiple metals significantly increased in concentration downstream of the town of Calnali when individually tested, including copper, lead, cadmium, aluminum, and zinc (Kruskal Wallis *P* < 0.001; Figures 6E, S3). On the other hand, other ions with known effects on fish neurobiology or olfaction showed similar or higher concentration in the Conzintla than in the Calnali downstream sites, including manganese, arsenic, and calcium (Figure S3). We also observed some seasonal fluctuations in metal concentrations, though we lack sufficient sampling to formally analyze these effects (Figure S5).

Ion concentrations were orders of magnitude higher in whole swordtail carcasses than in water (tissue:water ratio range 42–11289, median 430; Figure S6B), consistent with bioaccumulation of all metals measured. However, correlations between concentrations in water and tissue were not significant after controlling for the mean concentration of each metal across sites (ρ = -0.115, S = 58407*, P* = 0.3514). Moreover, PCA showed metal profiles varying upstream and downstream of the town of Calnali as in surface water, but loadings on the first two PCs showed discordant patterns for some ions (Figure S6C). Redundancy analysis found that site, month, and fish length had significant effects on fish’s ionic profiles (site *P* = 0.001, month *P* = 0.008, fish length *P* = 0.032, overall model *P* = 0.001, adjusted *R*^2^ = 0.302). More variance was associated with site (30.3% of total variance) than with month (4.1%) or fish length (3.2%). In comparison, redundancy analysis of water chemistry data also found a strong relationship between explanatory variables and ion profiles (site *P* = 0.001, month *P* = 0.001, overall model *P* = 0.001, adjusted *R*^2^ = .491). However, sampling month captured a much larger portion of the variance (23.3%) relative to site (36.9%). Metal bioaccumulation at the whole-organism scale thus seems at least partially decoupled from water chemistry.

### Olfactory disruption as a function of water chemistry

Our chemistry measurements showed that fish in downstream sites on the Río Calnali experienced elevated levels of a range of compounds that may impact olfactory function. Given limited evidence for gross bioaccumulation, we focused on potential effects of water chemistry on the sensory epithelial surface. Specifically, we evaluated evidence for histological changes in the olfactory rosettes of wild-caught fish from an upstream site, Aguazarca, and a downstream site, Calnali Low. H&E staining allowed us to observe the motile and sensory cilia which may be lost in the face of chemical exposure^55–57^. We found marginally lower proportions of ciliated surface area on the olfactory epithelium at Calnali Low (Mann-Whitney Test *P* = 0.08; Figure 7C). Likewise, AB/PAS staining clearly distinguished goblet cells through the deep blue staining of the protective mucoproteins they produce, and revealed significantly higher densities of goblet cells at Calnali Low (Mann-Whitney Test *P* = 0.003; Figure 7A–B, 7D), consistent with a compensatory response to shield the olfactory epithelium from toxicants. In contrast to the differences observed in the wild, 96 hours of laboratory exposure to ecologically relevant levels of ammonia, copper, humic acid, or their combination yielded no such differences in ciliary surface (Kruskal-Wallis *P* = 0.166) or goblet cell density (Kruskal-Wallis *P* = 0.406) in *X. birchmanni* (Figure 7E–F). Both histological metrics were comparable across all laboratory treatments to the values observed in the upstream Aguazarca population (Figure 7C–F).

**Figure 7.**
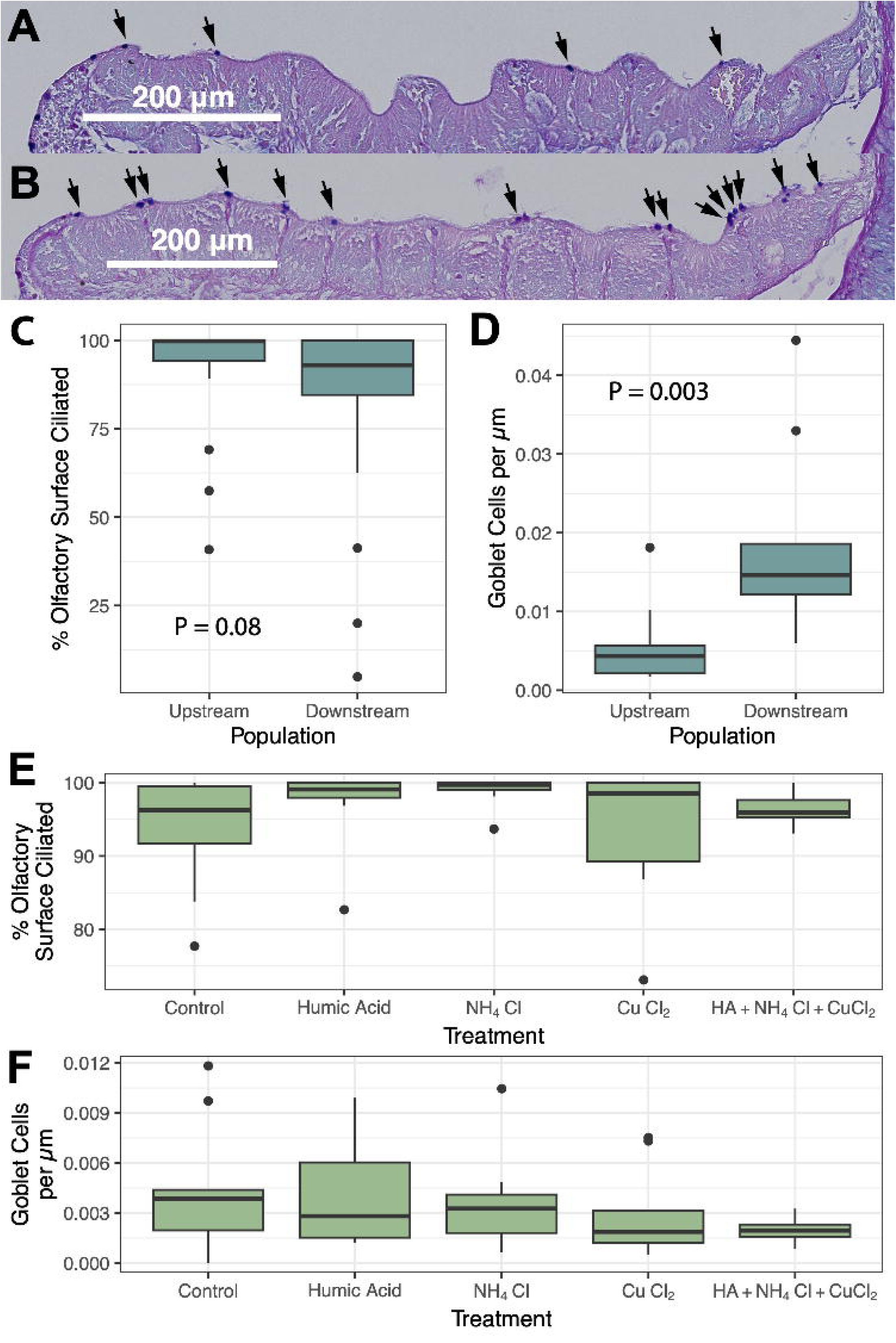
Histological phenotypes in wild-caught and laboratory-exposed fishes. **A–B)** Representative AB/PAS stains of olfactory rosettes of *X. malinche × birchmanni* hybrids from sites **A)** upstream (Aguazarca), and **B)** downstream (Calnali Low) of the town of Calnali. Mucoproteins are stained blue by AB/PAS, eosin counterstain appears as pink. Arrowheads indicate locations of goblet cells, with the pictured individuals exhibiting the median count for each site. **C–D)** Olfactory histological measurements. Thick lines denote group medians, boxes show interquartile ranges (IQR), and whiskers and points show samples inside and outside 1.5 × IQR, respectively. **C)** Proportion of each fish’s sensory epithelium with visible cilia, visualized with H&E staining (upstream *n =* 27, downstream *n* = 29). **D)** Number of mucus-producing goblet cells per µm of olfactory rosette surface area, visualized with AB/PAS staining (*n =* 9 individuals each). *P*-values indicate results of Mann-Whitney *U* Test for difference between sites. **E–F)** Results from laboratory chemical exposure experiment (*n* = 10 individuals each). Adult *X. birchmanni* were exposed to elevated dissolved organic matter, ammonia, or copper (delivered as humic acid, NH_4_Cl, and CuCl_2_, respectively) or all the above for 96 hours, and compared to control fish with no exposure. **E)** Ciliated surface and **F)** goblet cell density were measured as in **C–D**. Note difference in Y-axis scale from **C–D**. See Figure S7.

## Discussion

With a fine-scale view of the genetic and environmental landscape of hybrid zones, we can move past a binary picture of hybridization’s occurrence to assess the specific changes and mechanisms driving anthropogenic hybridization. Here, we explore spatial covariation in population structure, assortative mating, and environment as a proxy for the changes that may have driven initial hybridization between *X. birchmanni* and *X. malinche*. We then use targeted analyses within the Río Calnali to test the plausibility of one hypothesized driver: the disruption of olfactory-based mate choice by chemical pollution from human development. Although our results are primarily based on correlations, they are consistent with this hypothesis, and lay the foundations for further manipulative studies.

By surveying genome-wide ancestry distributions across dozens of sites, we documented the outcomes of the complex interactions of migration, ecological selection, and mate choice in structuring hybrid zones. Migration influenced ancestry structure in all drainages, with waterfalls and rapids isolating near-pure *X. malinche* populations upstream in the Ríos Pochula and Huazalingo, and ancestry distributions shifting on either side of subterranean stream reaches in the Ríos Calnali and Conzintla (Figures 1, 3). Though we found some degree of gene flow between *X. birchmanni* and *X. malinche* in all drainages, the amount and spatial distribution of admixture differed starkly between streams (Figure 3). At one extreme, the Río Conzintla showed two ancestry clusters with limited admixture across multiple sites. The functional consequences of this restricted admixture may be limited by genome-wide selection against introgression^43^, and populations with similar admixture rates have been identified as warranting conservation in other anthropogenic hybrid zones^58^. The stable Conzintla hybrid zone thus serves as a contrast to other streams, and environmental comparisons may illuminate the potential drivers of breakdowns in assortative mating.

The Río Huazalingo and Río Pochula were both intermediate in water chemistry to the Río Conzintla and the Río Calnali, but showed divergent ancestry patterns. The Huazalingo, like the Conzintla, had a zone of sympatry between two distinct ancestry clusters, but in the Huazalingo this zone was temporally unstable (Figure S1A), lower in elevation (769 vs. 1020 m; Table S1), and limited to the area immediately downstream of a waterfall. This pattern could be explained by a combination of migration barriers and ecological selection, with *X. birchmanni*-like individuals prevented from migrating upstream by the waterfall, and *X. malinche*-like individuals outcompeted when passing below it. In contrast, the Río Pochula hosted near-pure *X. malinche* at even lower elevations (522 m; Table S1), but no sites with bimodal ancestry structure. Rather, this stream appeared more similar to classic clinal hybrid zones, where ancestry patterns are shaped by the countervailing effects of gene flow and selection against hybrids^59^.

Why might rivers with similar water chemistry differ in population structure? Potential explanations might hinge on differences in watershed area and thus stream volume (Figure S4A) at the point of interspecific contact. The attendant differences in swordtail habitat availability, population densities, and signaling environment might alter pre-mating isolation, as both demography^7^ and individual condition^60^ are known to affect hybridization rates. Alternatively, past anthropogenic impacts in the Pochula may have caused a collapse of population structure, which persisted beyond the disturbance due to a loss of ancestry-assortative mating in admixed individuals^61–63^. The diversity of population structures we observed across watersheds is unsurprising given the multitude of factors known to affect mate choice and hybrid fitness^6–11,13,24^. That said, similar surveys of additional streams may provide the necessary contrasts to distinguish among some potential drivers.

The dramatic changes in ancestry structure in the Río Calnali from upstream to downstream allowed us to ask more direct questions about the relationship between environmental factors and population structure. We observed strong bimodality in genome-wide ancestry at sites upstream of the town of Calnali, with individuals falling into an *X. malinche-*like ancestry cluster and an *X. birchmanni*-like cluster. These results align with prior work documenting the presence of strong assortative mating by ancestry within ancestry clusters at the Aguazarca site on the Río Calnali^52^. In the less than two kilometers where the stream traverses the town of Calnali, this bimodal population structure collapsed into a hybrid swarm, with admixture proportions spanning those of the upstream clusters (Figure 3).

Likewise, mother-embryo sequencing and simulations demonstrated that assortative mating is substantially weakened, but not entirely absent, at downstream sites on the Río Calnali (Figure 4). Ongoing assortative mating could be driven by some combination of downstream migration of pregnant mothers, variable impacts of pollution on different sensory pathways (e.g. visual vs. olfactory cues)^64,65^, or fluctuations in sensory disruption as a result of variation in environmental conditions (Figure S5). Regardless of the precise drivers, our findings support theoretical predictions that even modest reductions in the strength of assortative mating can result in the emergence of hybrid swarms^26^. The Río Calnali thus provides an opportunity to study the environmental determinants of reproductive isolation through comparisons within and between streams.

We next confirmed that anthropogenic impacts vary over space within our study area. As expected, the Calnali drainage was unique both in the scale of built infrastructure relative to catchment area, as well as the proximity of development to the stream (Figure 5). Runoff from agricultural lands could also contribute to the chemical and biological variation we observed, but may be a less likely explanation given the relatively small differences in agricultural land area within and between drainages (Figure S2). We then identified stream-specific water chemistry signatures, which we speculate may be driven by geology and mining impacts in the Conzintla drainage (Figure S4B), as well as orthogonal chemical changes between sites on the Río Calnali above and below the town of Calnali (Figure 6). Both agricultural and urban development have been linked to some of the changes in water chemistry that we observe in the downstream Río Calnali^66–70^. In particular, many metals whose concentrations change along the course of the Río Calnali are associated with both domestic wastewater and runoff from roads^68^.

Among the compounds that varied spatially, a substantial number have been previously shown to affect fish olfaction^19,71,72^ or neurobiology^73–75^. Many of these parameters were especially elevated in the areas downstream of the town of Calnali, and thus are potential candidates for the drivers of ancestry structure breakdown in this stream. Turbidity also increased in downstream Calnali sites, and though interspecific mate discrimination in these species is thought to be primarily olfactory^76,77^, a disruption of visual preference could nonetheless contribute to hybridization^78^. In contrast, some ions with documented olfactory or neurological effects were most concentrated in the Conzintla (Figure S3)^23,79–81^. This highlights the need both for ecotoxicological studies in *Xiphophorus* to establish species-specific sensitivity (i.e. EC_50_ values), and for careful consideration of how the total chemical environment may modify the effects of individual compounds. For example, variation in pH may generate interaction effects, as low pH increases metal bioavailability and toxicity (Figure S3)^71^.

We next tested for metal accumulation in whole swordtail bodies, and found mixed evidence. Bioaccumulation was present but modest compared to levels previously reported after intentional ecotoxicological exposure^82^, and varied less over time than concentrations in the surrounding water (Figures S5–S6). The lack of correlation between relative concentrations of ions in water and in aggregate tissue implies that any bioaccumulation was not mediated by direct absorption from water (i.e. bioconcentration) alone. This decoupling may be driven by site-wise differences in fish morphology and tissue allometry (Figure S6A), differential exposure from diet^83^ or differential sequestration in varying habitat substrates^84^. Regardless, our results did not support gross metal toxicity as a trigger for hybridization in the Río Calnali. Anthropogenically introduced metals might still affect reproductive isolation through tissue- and ion-specific accumulation (e.g. in gonads, brain, sensory organs) or through direct effects on extracellular interactions between sensory epithelia and signaling molecules. Given prior work on sensory alterations by humic acid^22,85^ and metals^84^, we identified the latter mechanism as a promising candidate for targeted observations.

Having established differences in both the degree of population structure and in chemical environment between sites in the Río Calnali, we explored the loss of olfaction-based assortative mate preferences as a mechanism which might connect the two. Our histological analysis of fishes in the upstream and downstream Calnali showed changes in morphology downstream consistent with olfactory damage (Figure 7A–D). The loss of epithelial cilia has previously been observed after exposure to metals and anthropogenic hydrocarbons^86,87^, and could directly reduce olfactory sensitivity by disrupting water flow through the nares^88,89^ or the function of olfactory sensory neurons^90^. Likewise, increased goblet cell densities have been linked to nasal irritation in other systems, and may protect epithelia from noxious chemical exposure^91–93^. By showing changes in olfactory structure *in situ*, this work complements prior *ex situ* experiments showing that water chemistry manipulations can alter assortative olfactory preferences in *X. birchmanni*^19,23^.

However, our short-term laboratory exposures of *X. birchmanni* to ecologically relevant levels of ammonia, copper, humic acid, or their combination did not recreate the histological differences seen *in situ* (Figure 7E–F). The effects seen in wild-collected fish may thus be dependent on the cumulative and interactive effects of numerous chemicals, exposure for longer periods, or exposure at a more sensitive developmental stage^94–96^. Definitively linking the observed histological differences to the chemical environment, rather than a covarying biotic factor (e.g. pathogen exposure)^97^, will require additional evidence from inter-stream comparisons and experiments. Likewise, whether these histological changes themselves weaken assortative mating, or are correlated with other causative changes (e.g. neural accumulation of toxicants^98^, water chemistry effects on signal transduction^22^, individual condition^60,99^, or conspecific encounter rates^7^) is an exciting direction for future work.

Taken as a whole, our study presents an integrative window into the environmental and organismal drivers of hybridization between *X. birchmanni* and *X. malinche*. By showing that water chemistry differences coincided with changes in population structure, we support the relevance of prior behavioral work demonstrating the disruption of assortative mating in these species following water chemistry changes^19,23^. It remains to be seen whether reproductive isolation might be re-established between these species through a combination of habitat restoration, immigration, and natural selection against hybrids, as has been observed in other cases of anthropogenic hybridization^100^. Though many studies have documented a broad connection between environmental conditions and hybridization in freshwater fishes^5,24^, our analyses of assortative mating and olfactory histology provide a deeper understanding of proximal mechanisms by which water chemistry changes might disrupt reproductive isolation on small spatiotemporal scales. Further characterizing the drivers of hybridization will both expand our understanding of the processes which generate biodiversity, and enable improved management of unintended anthropogenic impacts.

## Resource Availability

### Lead contact

Further information and requests for resources and reagents should be directed to and will be fulfilled by the lead contact, Molly Schumer.

### Materials availability

This study did not generate new unique reagents.

### Data and code availability

- Raw sequencing reads used in this project are available under NCBI SRA Bioprojects PRJNA1346846, PRJNA1305547, PRJNA744894, PRJNA930165, and PRJNA610049.
- All other datasets necessary to recreate the results of this publication are available on Dryad (https://doi.org/10.5061/dryad.nvx0k6f5h).
- All original code used to generate results are available as of the date of publication on Zenodo at https://doi.org/10.5281/zenodo.19637379 and GitHub at https://github.com/Schumerlab/Lab_shared_scripts.
- Any additional information required to reanalyze the data reported in this paper is available from the lead contact upon request.

## Acknowledgements

The authors thank the Schumer and Rosenthal laboratories for helpful feedback on earlier manuscript versions. Stanford University and the Stanford Research Computing Center provided computational support for this project. We thank Dra. Susana Saval Bohórquez of the UNAM Environmental Engineering Laboratory and Dr. Guangchao Li and Douglas Turner of the Stanford Environmental Measurements Facility for experimental support. We are deeply grateful to the delegates and landowners of the Calnali region for their engagement in our work and for sharing access to their communities’ waterways. This work was supported by a Knight–Hennessy Scholars fellowship and NSF GRFP 2019273798 to B.M.M., a CONACyT fellowship to G.I.J., NSF Awards 1723266, 1755327 and 1354172 and a grant from the Fondazione Cariparo to G.G.R., NIH grant 1R35GM133774 to M.S., and Human Frontiers in Science Grant RGY0081 to M.S. & C.M.R.

## Author Contributions

Conceptualization, B.M.M., C.M.R., M.S., and G.G.R.; Investigation, B.M.M., W.F.R.-D., D.L.P., T.T.L.Y., T.R.G., G.I.J.-R., C.Y.P., E.N.K.I., G.A.P., S.M.B., A.E.D., R.S., and J.J.B.; Software, B.M.M. and M.S.; Formal analysis, B.M.M.; Resources, C.M.R., M.S., and G.G.R.; Writing original draft, B.M.M. and M.S.; Writing — review and editing, B.M.M., W.F.R.-D., D.L.P., G.I.J.-R., E.N.K.I, G.A.P., S.M.B., G.M.V.A., C.G.-R., C.M.R, M.S., and G.G.R.; Supervision, G.M.V.A. and C.G.-R.; Funding acquisition, C.M.R., M.S., and G.G.R.; Project administration, G.M.V.A.

## STAR Methods

### Key Resources Table

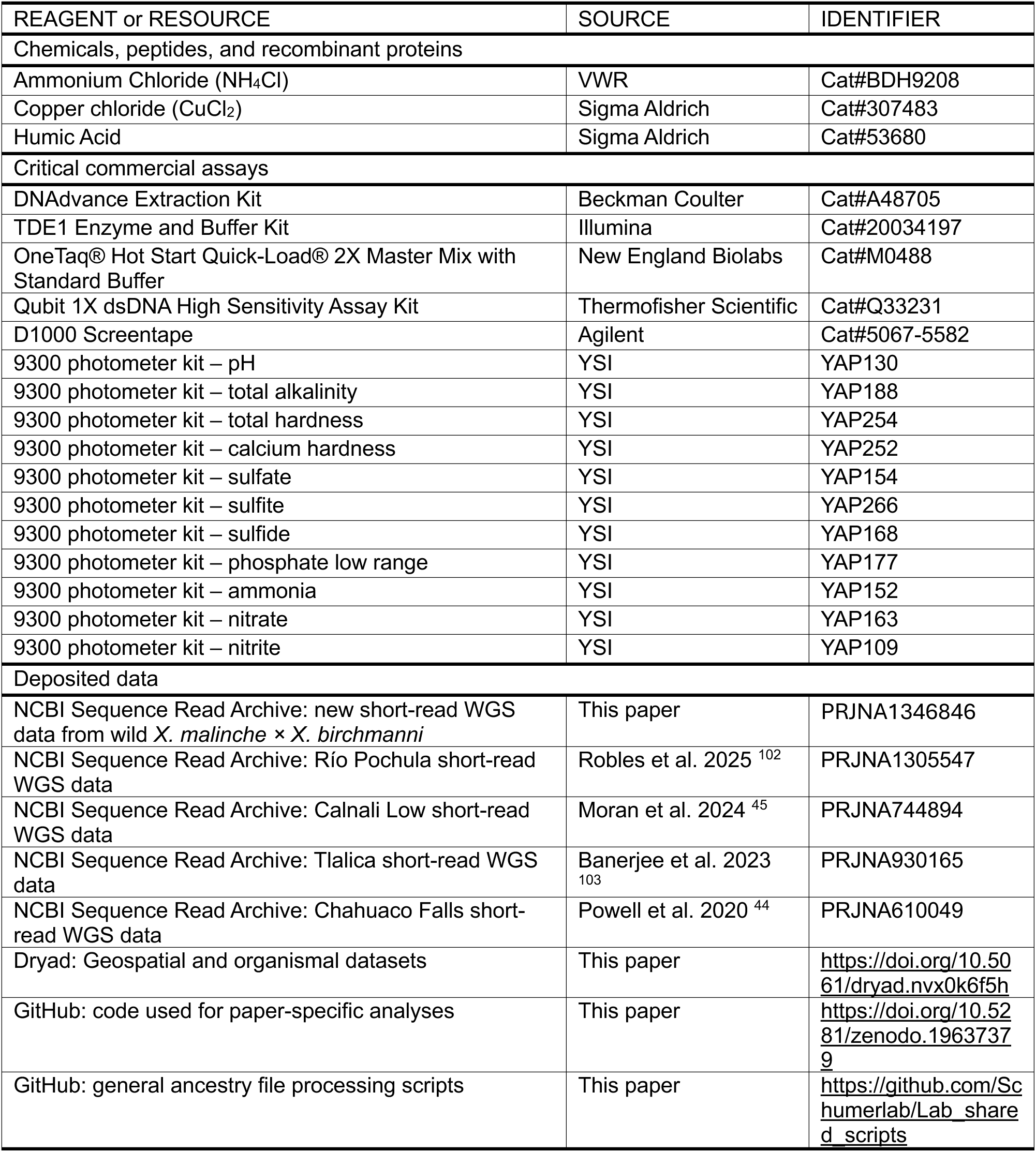
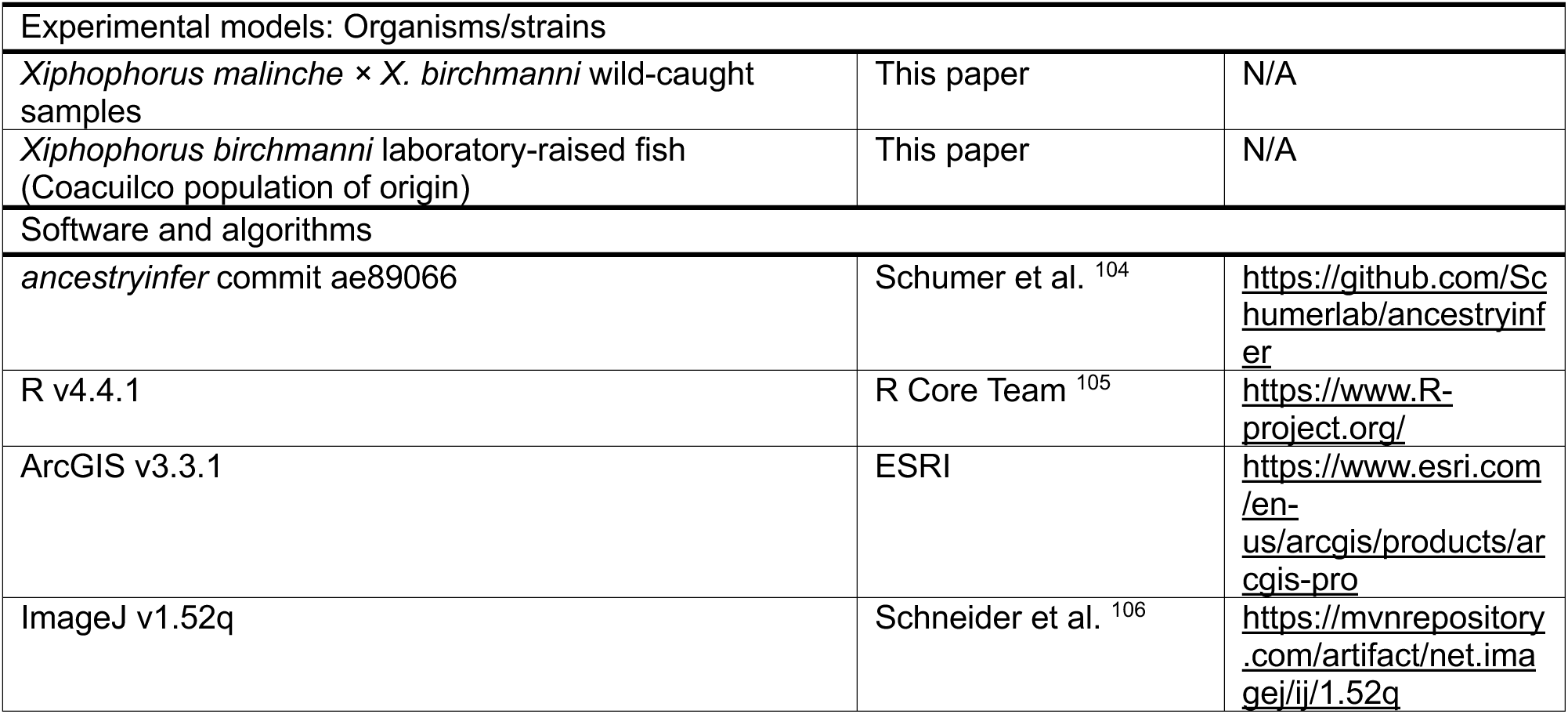

### Experimental model and study participant details

*Xiphophorus* fishes were collected at 48 sites across four stream basins, which together compose the Río Atlapexco drainage of Hidalgo, Mexico (see Table S1 for full list with GPS coordinates, and Table S2 for sample size by year), with permission from the Mexican government (Permiso de Pesca de Fomento no. PPF/DGOPA-064/20) and Stanford University animal welfare protocols (Stanford APLAC protocol #33071). Fish were collected using baited minnow traps and anesthetized in a buffered solution of 100 mg/mL MS-222 dissolved in stream water. Fin clips were taken and stored in 95% ethanol for low-coverage whole-genome sequencing, after which fish were allowed to recover then returned to the pool from which they had been sampled. A subset of female fish from the Calnali Mid (*n* = 22) and Calnali Low (*n* = 74) sites (Figure 1) were euthanized and stored in 95% ethanol for later dissection of embryos and analysis of assortative mating (see below). In addition, 30 fish each from the Aguazarca and Calnali Low sites were decapitated following euthanasia in May and September 2022, with whole heads preserved in 10% neutral buffered formalin for olfactory histology (see below). Finally, six fish each were collected for whole-body ICP-MS from CAPASC Fuente de Agua, Aguazarca, Calnali Low, and Tlalica in November 2021 and February 2022 (total *n* = 12 individuals per site), with the length and weight of each fish measured before euthanasia and storage at -80°C. *X. birchmanni* used in laboratory chemical trials were born and raised in the Stanford fish facility (source population Río Coacuilco, 21°5’51.16”N 98°35’20.10”W). Fish were late juveniles or adults (>4 months old) before the beginning of experimentation in July and August 2023. Experimental trial groups were balanced by sex, with 2 females and 3 males per treatment in July, and 3 females and 2 males per treatment in August. Fish were housed in groups of 5 in 10-gallon tanks on a 12/12 light dark cycle at 22°C and fed twice daily. All individuals were healthy and had not been involved in previous procedures.

### Method Details

#### Low-coverage whole-genome sequencing

DNA was extracted from preserved fin clips using the Agilent DNAdvance extraction kit (Beckman Coulter, Brea, California). The protocol followed the manufacturer’s instructions for tissue extraction except that we used halved volumes in all reactions. Extracted DNA was quantified on a BioTek Cytation5 (Agilent, Santa Clara, CA) microplate reader and diluted to 2.5 ng/µl for library preparation. We used a previously described tagmentation-based protocol to prepare DNA libraries for low-coverage sequencing^41^. We prepared a tagmentation master mix to enzymatically shear diluted DNA using 0.25 µl of TDE1 Tagment DNA enzyme (Illumina no. 20034197), 1.375 µl of TD1 buffer (Illumina no. 20034197), and 1.375 µl of Tris-HCl MgCl_2_ buffer (20 mM Tris-HCl, 10 mM MgCl2) per sample. We added 3 µl of tagmentation master mix to 2 µl of diluted DNA and incubated at 55°C for 5 min. We prepared a PCR master mix with per-sample volumes of 8.9 µl of OneTaq Hot Start Quick-Load 2X Master Mix (NEB no. M0488) 3.5 µl of nuclease-free water, and 1.2 µl of 10 µM i5 index. We combined 5 µl of tagmented DNA, 1.5 µl of 10 µM i7 index, and 13.5 µl of the PCR master mix per sample and amplified the PCR reaction for 12 cycles. We used custom i7 indexes. PCR products were pooled and purified using 18% SPRI beads. Libraries were quantified on a Qubit fluorometer (Thermo Scientific, Wilmington, DE), and library size distribution was visualized on an Agilent 4200 Tapestation (Agilent, Santa Clara, CA). Sequencing was carried out at Weill Cornell Medical College, Harvard Bauer Core, and Admera Health Services (South Plainfield, NJ) on an Illumina HiSeq 2500 (*n* = 526), HiSeq 4000 (*n* = 1289), NovaSeq X (*n* = 221), and NovaSeq X Plus (*n* = 709). Library formats included 150 bp paired-end reads (*n* = 660), 75 bp paired-end (*n* = 691), 100 bp single-read (*n* = 182), and 100 bp paired-end (*n* = 276). The average nominal coverage (base pairs sequenced / *X. birchmanni* genome size) of individuals included post-filtering was 0.65X. Individual-specific information is available in the metadata of the relevant NCBI BioProjects (see Data Availability statement).

#### Water chemistry surveys

We chose to focus water quality sampling on a subset of 32 sampling sites that captured the full range of variation in ancestry structure present in each stream and collected measurements from these sites between February 2022 and September 2024 (Figure 1). Prior work has shown that reproduction in *Xiphophorus* is seasonal, with pregnancy rates peaking in early summer in both *X. birchmanni* and *X. malinche*, and reproduction nearly nonexistent in high-elevation populations during the fall and winter^99,107^. We therefore focused our studies of water quality on the region’s dry season (February through early June), under the hypothesis that water chemistry in the period coinciding with reproduction would have the most direct effect on mate choice and species barriers. However, we also conducted limited sampling in September during the rainy season (July through November) to understand how water chemistry might vary throughout the year, and how this might affect our conclusions about the link between water chemistry and population structure.

Due to changes in equipment availability over time, sample sizes for individual water quality measurements varied between 0 and 13 per location and measure; for details on all measures collected and sample sizes per location, see Data S1A–B. All water chemistry measurements were taken from streambanks or exposed rocks to minimize disturbance of bottom sediments, and samples were collected 2–10 cm below the stream surface with laboratory-grade plastic bottles, 50 mL Falcon tubes and 50 mL sterile syringes. We measured temperature, conductivity, dissolved oxygen, and fluorescent dissolved organic matter (fDOM) using YSI EXO 2 and EXO 1s multiparameter sondes (Xylem Inc.) held 2–10 cm below the stream surface. Turbidity was measured in nephelometric turbidity units (NTUs) with an Orion AQ4500 Turbidity Meter (Thermo Fisher). Using a YSI 9500 photometer, we applied colorimetric assays to measure pH, total alkalinity, total hardness, calcium hardness, and concentrations of sulfate, sulfite, sulfide, phosphate, ammonia, nitrite, and nitrate. Turbidity measurements and colorimetry tests were performed at the site of sampling to ensure the accurate measurement of unstable chemicals, while further measurements were performed on collected water samples. For measurements of dissolved organic carbon (DOC) and total nitrogen, we stored samples in glass vials capped with PTFE/silicone septa, which were combusted at 500°C for 1 hour and rinsed ten times with ultrapure water (Milli-Q^®^ Merck) prior to sample collection. Water samples were processed on-site by syringe-filtering through a 0.45 μm polyethersulfone (PES) membrane, acidifying to pH 2–3 using concentrated hydrochloric acid, and capping with no air bubbles remaining in the vial. DOC and N measurements were carried out on a Shimadzu TOC-L analyzer using the TC/IC and TN protocols, respectively, at the Environmental Engineering Laboratory of the National Autonomous University of Mexico (UNAM) and the Environmental Measurements Facility at Stanford University. Total and dissolved metals were measured from samples collected in 50 mL Falcon tubes triple-rinsed with sample water, with samples for dissolved metals filtered through 0.45 μm PES and all samples acidified to a pH of ∼2 using trace metal grade nitric acid. Inductively coupled plasma mass spectrometry (ICP-MS) detection of 39 ions was carried out following a modified version of the APHA3030B/6020A methods by ALS Environmental (Waterloo, ON; see Data S1 for full list of ions, sample sizes, and quality control). Given that many metals in the filtered samples were below the lower detection limits of ICP-MS, we used ICP-MS measurements of total concentration from unfiltered samples in all downstream statistical analyses.

#### ICP-MS analysis of fish tissue in the Río Calnali

To understand how changes in concentration of metals in rivers might lead to differential bioaccumulation in swordtails, we measured ion concentrations in whole fish at a subset of our sampling sites on the Río Calnali. In November 2021 and February 2022, six fish from CAPASC Fuente de Agua (CAPAC; #15 in Figure 1 and Table S1), Aguazarca (AGZC), Calnali Low (CALL), and Tlalica (CAPS; #25 in Figure 1 and Table S1) were measured for length and weight before euthanasia, storage at -80°C, and submission to ALS Environmental Services for ICP-MS analysis. At ALS, whole fish were homogenized, subsampled, and microwave digested (EPA Method 3052) before collision-reaction cell ICP-MS (EPA Method 6020B). Where sufficient tissue mass was present, moisture content was also estimated for each sample as the proportional weight loss of a sample after drying at 105°C. Given that many samples were below the weight necessary for moisture analysis, all downstream analyses were performed on a wet-weight basis.

#### Histology in wild-caught and laboratory fish

Like their relatives^108^, *X. malinche* × *birchmanni* hybrids have plate-like olfactory rosettes with beds of sensory epithelia surrounded by ridges of nonsensory tissue (Figure S7A). Sensory epithelial beds contain a combination of ciliated and microvillous olfactory neurons, which collect and transmit chemical information to the olfactory bulb, and ciliated sustentacular cells, which increase flow over the beds to heighten olfactory sensitivity^86,109^. The intervening nonsensory ridges are composed largely of connective tissue and goblet cells, the latter of which are responsible for producing the glycoprotein-based mucus which physically shields the epithelium from irritants^91,108^. Based on reports from the literature^55–57,91,92^, we quantified the loss of cilia from the sensory epithelial surface and the proliferation of goblet cells in the nonsensory epithelium and treated these two measures as a proxy for olfactory disruption.

To understand whether polluted water was directly impacting the sensory periphery of fish in *X. malinche* × *birchmanni* populations, we collected a total of 30 fish each from the Aguazarca and Calnali Low populations in May (*n =* 10 per site) and September (*n =* 20) 2022. Fishes were euthanized by MS-222 overdose and decapitation, after which heads were fixed in 10% buffered formalin for at least 48 hours. Fixed crania were dissected to remove the gills and lower jaw, decalcified for 8 hours in 1X PBS with 10% formic acid, then dehydrated for 20 minutes each in 30%, 50%, and 70% ethanol before long-term storage in 70% ethanol. Samples were submitted to Histo-Tec Laboratories (Hayward, CA) for xylene clearing, paraffin embedding, sectioning, and staining. Horizontal sections were taken at 5 μm intervals with a rotary microtome after removing 75–150 μm of tissue from the dorsal surface, with deeper sectioning needed to reach the olfactory epithelia in larger individuals. One slide per individual was prepared with hematoxylin and eosin (H&E) staining to observe gross morphological changes (i.e. presence/absence of ciliary on olfactory epithelium). Samples from one collection (May 2022) were also subject to Alcian Blue / Periodic Acid-Schiff stain for mucoproteins to identify mucus-producing goblet cells, with eosin as a counterstain. Individuals for whom olfactory rosettes were not present in the Histo-Tec slides were excluded from further analysis (final cilia *n* = 27 for Aguazarca, *n* = 29 for Calnali Low; final goblet cell *n* = 9 individuals for both populations). Slides were scored by a single blinded observer at 1000X magnification using oil immersion. Using H&E slides, we defined the bounds of each sensory epithelial bed, and estimated the proportion of the surface area of each bed on which cilia were visible to the nearest 10%. From AB/PAS slides, we counted all cells positive for AB/PAS staining on the surface of each rosette. We also scanned and imaged each slide at 40x magnification via a Biotek Cytation5 microplate reader, utilizing the in-place ruler on the image to calculate the length of each bed and rosette in ImageJ.

Finally, we designed an experiment to test if laboratory exposure to ammonia, copper, and dissolved organic matter, alone or in combination, could replicate histological changes observed in the wild. We chose these three compounds as known neurological and olfactory toxicants in fishes^19,71,110^ which were elevated in the site of population structure collapse downstream of the town of Calnali (Figure 6). In each round of testing, adult *X. birchmanni* were exposed to either 0.25 mg/L total ammonia (delivered as NH_4_Cl, VWR BDH9208), 2 μg/L copper (delivered as CuCl_2_, Sigma Aldrich 307483), 20 mg/L humic acid (Sigma Aldrich 53680), all three treatments combined, or a no-treatment control. These concentrations were chosen to match those observed in polluted sites downstream of the town of Calnali (ammonia and copper) or to match prior studies showing effects on *Xiphophorus* mate choice (humic acid)^19^. Concentrations were maintained by daily 50% water changes, and water samples were taken before and after water changes to ground-truth concentrations in the tanks. After 96 hours of exposure, fish were euthanized by lethal MS-222 overdose and decapitation, after which heads were fixed in 10% buffered formalin for at least 48 hours. The experiment was run in two batches, in July and August 2023, for a total of 10 fish per treatment. Horizontal cranial sections were prepared, H&E and AB/PAS staining were performed, and ciliary coverage and goblet cell densities were quantified as described for wild-caught fish. Ground-truth water samples were tested for the concentration of ammonium, dissolved organic carbon, and total copper on a Westco SmartChem 200 Discrete Analyzer, a Shimadzu TOC/TN Analyzer (TOC-L), and a Thermo Scientific iCAP RQ ICP-MS, respectively. These tests confirmed that we achieved the desired change in ammonia and copper concentrations to mirror those seen between natural sites, though the change in dissolved organic carbon with our treatment of humic acid was lower than the range of natural variation (Figures S3D, S7C).

### Quantification and Statistical Analysis

#### Local ancestry inference

Hybrids between *X. birchmanni* and *X. malinche* harbor ancestry from both species that varies locally along the genome and globally between individuals and populations. We used an approach developed by our group for accurate local ancestry inference in *X. malinche × X. birchmanni* hybrids that relies on a hidden Markov model and known ancestry informative sites that distinguish *X. birchmanni* and *X. malinche* to infer local ancestry (*ancestryinfer*)^104^. Both simulation studies^104^ and analyses of populations of known ancestry have indicated that this approach has a low error rate (∼0.1% per ancestry informative site)^45,103^. Likewise, inference of local ancestry is robust to a number of technical challenges, including parameter misspecification^104^.

We inferred local ancestry using *ancestryinfer* from whole genome sequence data for all individuals for which we collected more than 300,000 reads from the sequencing approach described above, with a maximum of 2,000,000 reads aligned per individual for computational efficiency. This initial coverage cutoff was based on previous simulation studies^104^ and resulted in 12 to 702 individuals per site post-filtering (median 32; Table S3). *ancestryinfer* outputs posterior probabilities of ancestry for all informative sites along the genome. In the case of *X. malinche × X. birchmanni* hybrids, this results in estimates of ancestry at 729,167 ancestry informative sites across 24 chromosomes. Since we were primarily interested in accurately estimating global admixture proportion for individuals along a spatial cline, we converted posterior probabilities to ‘hard calls’ using a threshold of 0.9. For example, if an individual had a posterior probability greater than or equal to 0.9 for *X. malinche* ancestry at a given marker, we converted that marker to the *X. malinche* state. If both ancestries had a summed probability ≤0.1, the site was called as heterozygous, and if a site did not have a posterior probability ≥0.9 for any ancestry state, we converted the site to missing data (average missing data across all individuals = 13.1%).

#### Estimation of genome-wide ancestry proportion

We used ancestry hard calls to estimate the proportion of the genome derived from each parental species. We tested two approaches to estimate this proportion. First, we divide the number of alleles called for the *X. malinche* state at a posterior probability threshold of 0.9 divided by the total number of alleles called, with heterozygous sites counted as 1 allele for each parental species. Second, we calculated ancestry taking the distance between sites into account. To do so, we identified tracts of each individual’s genome called with high confidence as *X. malinche*, heterozygous, or *X. birchmanni* ancestry, then took the mean ancestry across all tracts weighted by tract length. Both methods were carried out with custom scripts in Linux bash, Perl, and R. These two methods of ancestry estimation are expected to produce the same result if the distance between AIMs is not correlated with genome-wide ancestry. To test this, we applied both methods to 73 hybrid individuals collected at the Plaza site (#22 in Figure 1 and Table S1) in the Río Calnali. We observed that the allele count estimates were on average slightly less variable between individuals than the ancestry tract method (Figure S8A). However, the absolute deviation was small in all cases, with a maximum difference between the two methods of 3.99% and a mean difference of 2.51%. We therefore used the allele count method to estimate ancestry fractions of all individuals in our genetic dataset.

Given the wide range of coverage of individual genomes in our dataset (300,000–2,000,000 reads aligned in *ancestryinfer*), we also sought to test how genome-wide ancestry estimates varied as a function of read counts and library type. To do so, we chose five individuals of varying genome-wide ancestry from our 150bp paired-end libraries, and five individuals from our 100bp single-end libraries, to represent the highest and lowest effective coverage we would expect with our 300,000 read cutoff. We then drew from each individual’s fastq files with seqtk sample to create files with between 25,000 and 2,000,000 reads (or all reads in the individual’s entire fastq file, if < 2,000,000). *ancestryinfer* was run in two batches, one for all paired-end data and one for all single-read data. We then calculated hybrid indices for each output file as in the main text, and visualized hybrid index as a function of read count for each individual. Our downsampling analysis showed that decreasing read depths consistently decreased the amount of minor parent ancestry detected (Figure S8B). However, changes in inferred ancestry in paired-end libraries were slight among read sets that surpassed our minimum coverage cutoff of 300,000 reads, with a maximum shift in ancestry between 300,000 reads and maximum coverage of 2.6%. As expected given the decreased overall coverage, we found that single-read libraries may be more sensitive to the impacts of low read count than paired-end libraries (Figure S8B). However, the single-end read sets that passed the read count threshold showed similarly low variation in genome-wide ancestry (1.2%) between 300,000 reads and maximum coverage. We thus concluded that variation in read depth is unlikely to affect the conclusions of our study.

#### Survey of ancestry distributions across populations

With these results in hand, we calculated and visualized the distribution of admixture proportions in populations along the Río Calnali, Río Pochula, Río Huazalingo, and Río Conzintla (Figure 3). As previous results had indicated the presence of bimodal ancestry structure in upstream populations of the Río Calnali, we also calculated Hartigan’s dip statistic^111^, which quantifies departures from a unimodal distribution, on samples from each collection site. Since we used this test across many collection sites, we applied a Bonferroni correction for the number of tests. We also tested for spatial stability of ancestry fractions in the downstream Rio Conzintla by applying a Kolmogorov-Smirnov test to all possible pairs of sites between the end of the zone of sympatry and the point at which the stream temporarily passed underground (sites 45–49 in Table S1).

We recognized that variation in sample sizes across sites (Table S3) might complicate our interpretation of these results, and so we tested the effects of sample size on our ability to detect significant ancestry structure (Table S4). Since our primary concern was that low sample sizes might reduce our power to detect significant departures from a unimodal ancestry distribution, we performed a down-sampling analysis and tested its effects on our inference of population structure. Specifically, we chose to match the sample size at Pezmatlán (*n* = 22), the smallest sample size in the Río Calnali, which was below the median sample size across sites (*n* = 32.5). For each of 12 sites with sample sizes ranging from 22 to the maximum of our study (Table S4), we drew a sample size of 22 individuals without replacement from the pool of all individuals. We then calculated Hartigan’s dip statistic (*D*) for each subsampled population, and recorded the *P*-value of Hartigan’s dip test for unimodality. For each population, we calculated the mean *D* across 1,000 replicate subsamples, as well as the proportion of the time that we identified significant deviations from unimodality.

This analysis showed that down-sampling reduced the variance in *D* measurements between different sites included in our dataset (σ^2^ = 0.005 for full-sample size *D*, 0.003 for mean down-sampled *D*). Our down-sampling analysis also suggested that our power to detect ancestry structure even at low sample sizes was dependent on the specific ancestry distribution of the population. While we could reject unimodality the overwhelming majority of the time in populations with equal frequencies of *X. birchmanni*-like and *X. malinche*-like hybrids (significance frequency = 0.976 for Aguazarca, 0.908 for Xochicoatlán), down-sampling had much greater effects in sites with greater within-cluster variance in ancestry and in sites with unequal representation of ancestry clusters (0.845 for Calnali Mid, 0.429 for Piloncillo, 0.285 for Mixtla). In contrast, we rarely or never detected a non-unimodal ancestry distribution when sampling from a population inferred to be unimodal based on our full dataset (minimum 0.0, maximum 0.001).

As an additional approach, we performed a simple Pearson’s correlation between the sample size of a site and the *D* statistic inferred from our full genetic dataset. We found that the relationship between this metrics was negative but not significant (ρ = -0.252, t = -1.71, P = 0.095), implying that any effect of sample size variation might be to inflate signals of multimodality at sites with low sample sizes. Given that the majority of the sites where we detected significant departures from unimodality had larger sample sizes (minimum 29, all others > 51) we took this as evidence that our results reflected true positives, rather than false positives from under-sampling a distribution of high variation.

Given the broad period over which genetic samples were collected (between 2006 and 2024, with the highest sampling density between 2014 and 2022; Table S2) and the uneven spatial distribution of sampling in some years, we also sought to ensure that any spatial differences in ancestry structure we observed were not due to changing ancestry over time. In particular, we wished to exclude the possibility that a hybrid population could appear as two structured ancestry clusters in one sampling event, and as a unimodal hybrid swarm in another. We separated the ancestry data we had collected at each site by collection year, excluding sampling locations and years with sample sizes < 10, conducted a Kolmogorov-Smirnov test between pairs of sampling years with a Bonferroni correction for multiple testing, and plotted the ancestry distributions over time for sites that showed a significant change (Figure S1A-C). Of the 15 populations where fish were sampled across more than one year, we saw a significant change in the ancestry distribution in three of the populations (Kolmogorov-Smirnov *P <* 0.003; Figure S1). All these sites show a shift in the frequency of parental clusters over time, rather than an appearance or disappearance of admixed individuals, and so we concluded that the spatial variation we observed was not explicable by uneven sampling across time.

#### Analysis of ancestry-assortative mating

Previous work has shown that the distinct ancestry clusters in the Río Calnali upstream of the town of Calnali are maintained by strong ancestry-assortative mating^52^. Note that in this prior work, samples from the nearby sites Aguazarca and Calnali Mid were pooled for analysis, but we analyzed them separately here (Figure 1B). In the current study, we identified the Calnali Low population as one of the first sampling sites where this genetic structure breaks down along the Río Calnali. As a result, we were interested in evaluating the strength of ancestry-assortative mating in this population and making comparisons to upstream populations.

To do so, we analyzed mother-embryo samples from the Calnali Low population, including previously published samples collected in 2020^45^, as well as newly sequenced samples from a 2017 collection, and compared them to mother-embryo pairs collected from populations upstream of the town of Calnali between 2008 and 2018. Because swordtails are livebearing fish that exhibit multiple paternity, when we collected pregnant *X. malinche × X. birchmanni* females we also collected ∼10 offspring from 2–3 fathers^112^. Although we cannot directly assay the paternal genotypes, we can use the ancestry of the offspring to infer the ancestry of the male that sired each embryo, since offspring will inherit approximately 50% of their genome from each parent. Thus, the difference between the mother’s genome-wide ancestry and the genome-wide ancestry of her offspring is a useful proxy for paternal ancestry.

We have previously used this approach to quantify ancestry-assortative mating in upstream sites on the Río Calnali^52^, and repeated this approach here. Specifically, we asked about the observed difference between maternal and offspring ancestry genome-wide in each site, compared to expectations under a scenario where females mate at random with respect to male ancestry. To generate a distribution of offspring ancestry expected under no assortative mating, we performed simulations in R (version 4.4.1). For each focal female in our dataset (*n* = 74 for Calnali Low, *n* = 22 for Calnali Mid), we randomly sampled a mate from our full dataset of adults from that site (*n* = 702 for Calnali Low, *n* = 79 for Calnali Mid). Next, we drew the ancestry for a single offspring from a normal distribution centered on the average of the maternal and paternal ancestry, and with variance drawn from that observed among siblings in our real dataset. Specifically, we calculated the variance in ancestry across 1,000 randomly drawn pairs of siblings from each site; to be conservative in our inference of lost assortative mating, we used the 0.05 quantile of this variance (4 × 10^-6^) for our simulations, which was the same across both sites. We repeated this procedure until we had simulated offspring for all the focal females. We generated 1,000 replicate simulations and compared the observed and simulated differences in mother-offspring admixture proportions in each site. Finally, we compared the mean of the difference in mother-offspring admixture proportions between the two populations using a two-sample *t*-test.

#### Inferring the strength of assortative mating

We found evidence that the observed mother-offspring ancestry patterns in the Calnali Low site differed subtly from patterns expected if mating were truly random with respect to ancestry (Figure 4A–B). However, this difference from random mating was much weaker than that inferred in the population upstream of the town of Calnali (Figure 4A–B). To quantitatively compare the strength of ancestry-assortative mating consistent with the observed data, we used the same approach previously used to quantify the strength of ancestry-assortative mating at upstream sites^52^.

Briefly, we used an approximate Bayesian computation approach in R (version 4.4.1) to evaluate what levels of ancestry-assortative mating were consistent with our observed data. For each simulation, we drew a strength of assortative mating *P_assortative_* ranging from 0 (random mating) to 1 (perfect assortative mating) from a random uniform distribution. For each female, we then randomly drew a mate from the population ancestry distribution. If the mate’s ancestry proportion differed from hers by less than 5%, the mate was accepted. If it did not, we used a Bernoulli trial with success probability 1 – *P_assortative_* to determine if the mate was accepted. If the mate was rejected, we repeated the mate-drawing procedure until the female was matched with a mate, then simulated offspring as described above. This was repeated until all females were matched with a mate. We repeated this simulation procedure for a total of 3,000,000 simulations.

We used a simple rejection sampling approach to select simulations that closely matched the empirical data. For each simulation, we calculated the mean and variance of the difference in ancestry between mothers and their offspring. If both summary statistics fell within 5% of the observed value for the site (Calnali Low mean 6.6%, variance 0.46%; Calnali Mid mean 2.5%, variance 0.26%), we accepted the simulation. This allowed us to estimate the range of values of assortative mating consistent with our data. We plotted the posterior distribution of accepted assortative mating strengths and identified the maximum *a posteriori* (MAP) estimate. To test for a difference in the strength of assortative mating between populations, we first evaluated whether the inferred strength of assortative mating in one site fell within the 95% credible interval of the other site, and vice versa. To investigate whether the distributions were significantly different, we randomly matched each of the 1073 accepted simulations from Calnali Mid with one of the 5720 accepted simulations from Calnali Low, sampled without replacement. We then calculated the difference in mating proportion between each pair, and tested whether the median difference was significantly different from 0 using a Wilcoxon signed rank test. The inferred biological effect size was quantified by the MAP and 95% CI of the difference in mating proportion.

#### Geospatial analyses

To complement our measurements of water chemistry, we measured land use in each stream catchment as a proxy for anthropogenic disturbance. Briefly, we used Google Earth Engine to access Sentinel-2 multispectral imagery from the month of May in the years 2020–2023 ^113^, removed images with > 20% cloud cover, masked pixels flagged as clouds, then took the median intensity for each band and pixel to construct a composite image with bands 11 (1610 nm, short-wave infrared, 20 m^2^ resolution), 8 (842 nm, near-infrared, 10 m^2^), and 2 (490 nm, blue, 10 m^2^). This spectral combination provides better contrast between built infrastructure, herbaceous, and woody vegetation than the visible spectrum^114^. All further analyses were performed in ArcGIS Pro (version 3.3.1) with the Spatial Analyst and Image Analyst extensions. We segmented the three-band raster file using the Segmentation tool, with a spectral detail of 18, a spatial detail of 19, and a minimum size per object of 20 pixels. Next, we selected four land use types for classification: water, forest (continuous tree cover, excluding plantation-style agriculture), non-forest vegetation (grassland, pasture, or cultivated crops, including plantation-style orchard), and developed land (buildings, mines, and roads). We included water surfaces (primarily to prevent their misclassification as other land use types), forest as the primary undisturbed land cover in the area, and non-forest vegetation as an additional form of human disturbance beyond hardened infrastructure. Using 3 m^2^-resolution imagery of the study area in March 2022 downloaded from Planet Labs ^101^, we identified 60 polygons of each land class to serve as training samples. Given the short time frame between the capture of our Sentinel and Planet Labs image sets, we assumed that there would be negligible change in the proportion of land classes. We then used the Classification tool to perform object-based land use classification using a support vector machine algorithm with maximum 500 samples per class. We visually inspected the resulting classification for pixels misclassified as developed land, and removed these pixels from the dataset prior to further analyses (0.8% of all pixels).

To delineate catchments for each sampling site, we downloaded a digital elevation model (DEM) of the state of Hidalgo with 15 m^2^ resolution from the Mexican National Institute for Statistics and Geography^115^. Starting with this DEM, we calculated flow direction and flow accumulation, snapped each site to the nearest pour point pixel, and calculated the land area which drained into the stream upstream of a given site (Figure S4A), using the Flow Direction, Flow Accumulation, Snap Pour Points, and Watershed tools, respectively. To identify the land area with greatest potential for runoff into waterways (i.e. greatest influence on water chemistry), we used the Raster Calculator tool. We converted all cells which received flow from at least 3000 uphill cells into a polyline object representing the relevant waterways in our study area (Figure S4A). We then used the Buffer tool to identify the land area within 500 m, 100 m, and 50 m of these waterways, and used the Clip Raster tool to identify the subset of classified land use pixels within each of these buffers. Using the Zonal Histogram tool, we summed the pixel counts of each land use category in each site’s subcatchment, then calculated the proportion of the total pixels per subcatchment composed of each land use type. Finally, we calculated cumulative land use within a site’s watershed by summing the land use categories within its own site-specific subcatchment and within the subcatchments of all upstream sites.

#### Water chemistry surveys

Unless otherwise noted, all further statistical analyses were performed in R (version 4.4.1) with RStudio^116^. We first explored overall differences in water chemistry between sites and drainages using Principal Component Analysis (PCA). Given differing sample sizes of ICP-MS measurement relative to other measures, we conducted two separate PCAs, one for ICP-MS metals measurement (*n* = 66 measurements) and one for all other data (*n* = 58). For water chemistry parameters other than metals, we included only measurements that were consistently collected throughout the length of the field study, including dissolved oxygen (mg/L), fDOM (quinine sulfate units), conductivity (μS/cm), total hardness (mg/L CaCO_3_ equivalent), total alkalinity (mg/L CaCO_3_ equivalent), pH, nitrite (mg/L N), ammonia (mg/L N), phosphate (mg/L PO_4_), sulfate (mg/L SO_4_), and turbidity (NTU). Of the 39 ions measured in the metal measurements, we retained 30 that had more than one measurement above the lower detection limits of ICP-MS. For the purposes of this analysis, metals measurements below the detection limit were set to halfway between zero and the lower detection limit. For example, for cadmium, the detection limit was 5 ng/L, and we set values below the detection limit to 2.5 ng/L. While this approach has some limitations^117,118^ compared to alternative approaches using imputation^119^, we could not apply imputation approaches to our data given the possibility of systematic differences in concentrations of many of the metals between streams. However, we explored the potential impacts of this analysis choice on our conclusions by repeating PCAs with the measurements below the detection limit set to the detection limit or zero, and found our results were qualitatively unchanged.

We also used the PCA analysis to qualitatively compare patterns within and across streams and identify chemicals of interest for further statistical analysis. Specifically, we chose chemicals that had negative loadings on both PC1 and PC2 in either the non-metal or metal PCA because we found that this approach clearly separated sites downstream on the Río Calnali from the others (Figure 6). From this set of parameters, we next identified those which had evidence for affecting fish neurobiology, olfaction, or mate choice in the literature. These parameters included DOM^19^, turbidity^120^, nitrite^121^, ammonia^73,110^, copper^55,122^, lead^123^, cadmium^74,124^, zinc^125^, and aluminum^126^. In the analysis of ICP-MS metals data, where the effects of samples below the detection limit prevented the use of parametric tests, we used Kruskal-Wallis tests to identify differences in concentration within and between drainages. For all other variables, we used ANOVA and Tukey’s HSD tests to confirm differences between the downstream Calnali sites and the other groups, testing for differences in the natural log-transformed response variables as a function of drainage, sampling month, and their interaction.

Finally, we used the same approach to explore differences in water chemistry over different seasons across the sampling period. We constructed PCAs using data from all seasons for non-metal and metal parameters as in the main analysis, but excluding sulfate measurements from non-metal analysis due to missing measurements in September (Figure S5). We also observed the change in individual parameters with river distance in the Río Calnali drainage (Figure S5), which was the only area in which we collected more than one sample per site in September (1–3 measurements per site for non-metal parameters, 1–2 for ICP-MS metals). We treat this analysis as preliminary, as we are unable to fully disentangle the effects of space and time without intensive sampling across watersheds under a variety of weather conditions.

#### Fish tissue ICP-MS analysis

To identify gross morphological differences that might influence bioaccumulation, we used a Type II ANOVA and Tukey’s HSD to test for differences in total length and condition factor (weight / total length) as a function of site, month, and their interaction. We also calculated site-wise mean concentrations of each ion and tested the correlation between the concentrations in surface water and tissue after controlling for differences in mean metal abundance across all sites. Specifically, we constructed two linear models, one model for measurements in water, and one for measurements from whole fish, each with the ion measured as the explanatory variable and the log-scaled site-wise mean concentration as the response variable. We then took the Spearman’s rank correlation of the residuals from the two models. We applied principal component analysis to visualize the tissue concentrations of ions that were consistently measured above the detection limit across all sites (Ag, Al, As, Ca, Cd, Co, Cr, Cu, Fe, K, Mg, Mn, Mo, Na, Pb, Se, Zn). Finally, we tested for differences in tissue metal profiles between groups using redundancy analysis (RDA), with a response matrix composed of all tissue ion concentrations included in PCA, and site, month, and fish length as explanatory variables. For direct comparison between tissue and water data, we conducted an RDA with a response matrix composed of the same ions measured in water at the same four sites, with site and month as explanatory variables.

#### Histological data

We weighted ciliated proportions by bed length to calculate the total proportion of the sensory epithelium with visible cilia in each individual, then used a Mann-Whitney *U* test to test for differences in ciliated surface proportion between fish from Aguazarca and Calnali Low. Likewise, we tested for a significant difference between sites in the number of goblet cells, corrected for rosette length, using Mann-Whitney *U* tests. Kolmogorov-Smirnov tests indicated that the ciliated surface proportions were not significantly different between the two sampling months (*P* = 0.481), and that there was no significant difference between the two populations or sampling months in the number of sensory beds (population *P* = 0.7951, month *P* = 0.5238), individual bed length (population *P* = 0.08173, month *P* = 0.3268), or total bed length (population *P* = 0.6684, month *P* = 0.2415). For laboratory chemical exposure trial data, we tested for significant differences between treatments in the rates of ciliary loss and goblet cell densities using non-parametric Kruskal-Wallis tests.

## Supplemental Information

**Document S1.** Figures S1–S8 and Tables S1–S5.

**Data S1. Metadata for water chemistry measurements. Related to Figure 6**. **A–B)** Sample sizes of each water chemistry parameter at each site, **A)** excluding measurements from September as in main text analysis, and **B)** including measurements from September as in Figure S5. Site codes match those in Table S1. **C)** Key matching ALS sample codes from subsequent panels to the site, date, and sample type (dissolved or total concentration) of the sample. **D–JJ)** Complete records from ICP-MS water chemistry analysis. Panels represent data from **D–H)** February 2022, **I–M)** May 2022, **N–S)** September 2022, **T–Y)** May 2023, **Z–DD)** February 2024, and **EE–JJ)** September 2024. **D, I, N, T, Z, EE)** Detailed per-sample analysis results. **E, J, O, U, AA, FF)** Technical duplicate results. **F, K, P, V, BB, GG)** analysis of quality control samples of known concentration. **G, L, Q, W, CC, W)** Details of employed methodology, including lab location and method reference number. **H, M, S, Y, DD, JJ)** Ion-specific detection limits. **R, X, II)** record of recommended hold time exceedance. (Provided as Excel file)

