## Supplementary Information for "Increased rates of hybridization in swordtails are associated with water pollution"

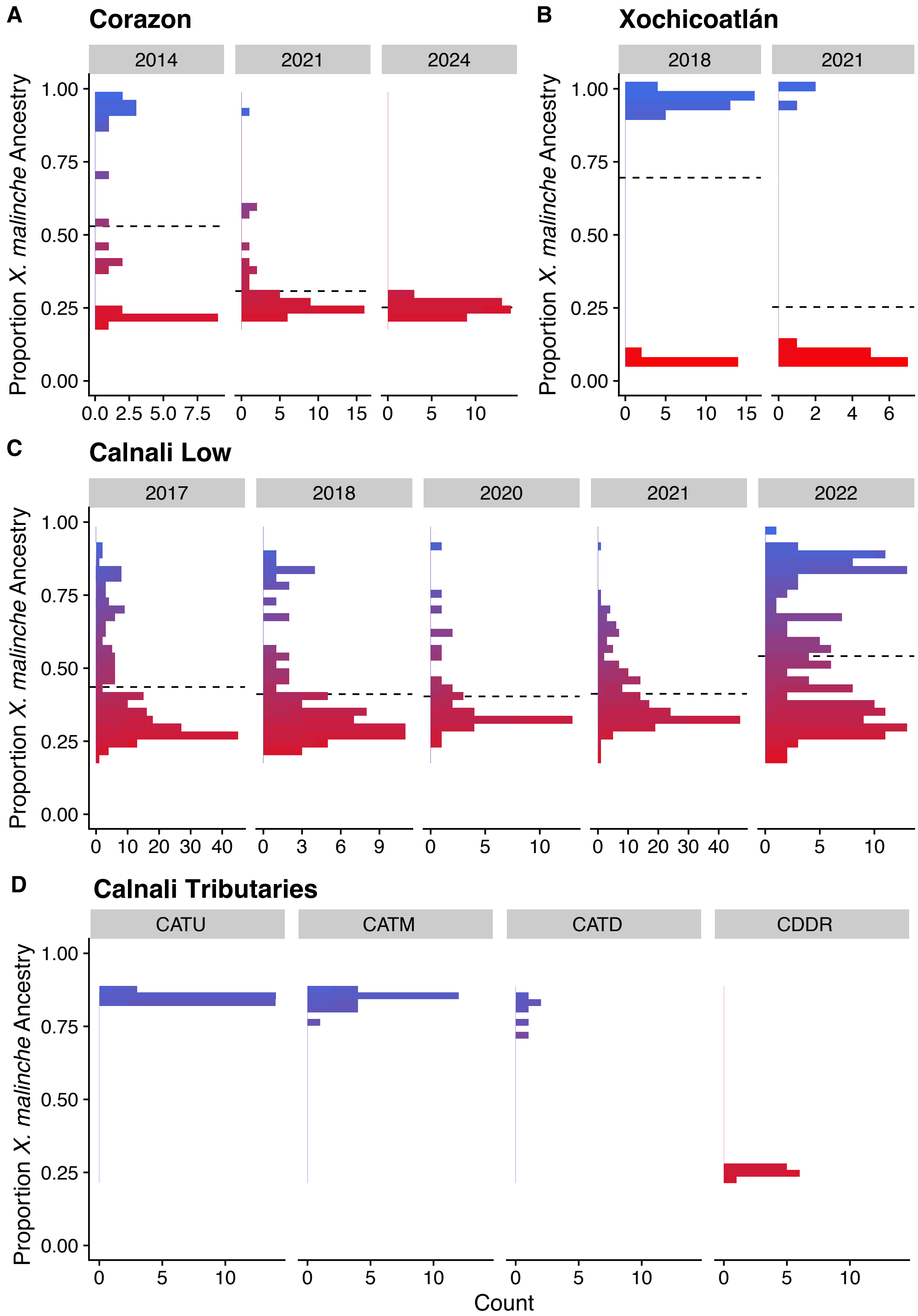


**Figure S1. Additional population ancestry distributions. Related to Figure 3. A–C)** Ancestry across sampling years at sites with significant differences in ancestry distribution over time (Kolmogorov-Smirnov $P < 0.003$). **A)** the Corazón site on the Río Huazalingo, **B)** the Xochicoatlán site on the Río Conzintla, and **C)** the Calnali Low site on the Río Calnali. The Y-axis denotes the proportion of the genome derived from the *X. malinche* parent (i.e. 1 is equivalent to 100% *X. malinche* ancestry genome-wide). The width of bars on the X-axis indicates the number of individuals in that ancestry bin. Based on Hartigan's dip statistic, we rejected a unimodal ancestry distribution at the Corazón site in 2018 (dip test $D = 0.132$, $P = 0.0002$), at the Xochicoatlán site in 2018 (dip test $D = 0.141$, $P < 1e\text{-}11$) and Calnali Low in 2022 (dip test $D = 0.066$, $P = 0.00002$). **D)** Ancestry distributions in four Río Calnali tributaries. Each tributary flows into the Río Calnali between the sites of Calnali Low and Tlalica. Sites are arranged with upstream sites on the left side of the X-axis, and downstream sites on the right. Abbreviations: CATU – Calnali Tributary Up, CATM – Calnali Tributary Mid, CATD – Calnali Tributary Down, CDDR – Ciudad de Ranas.

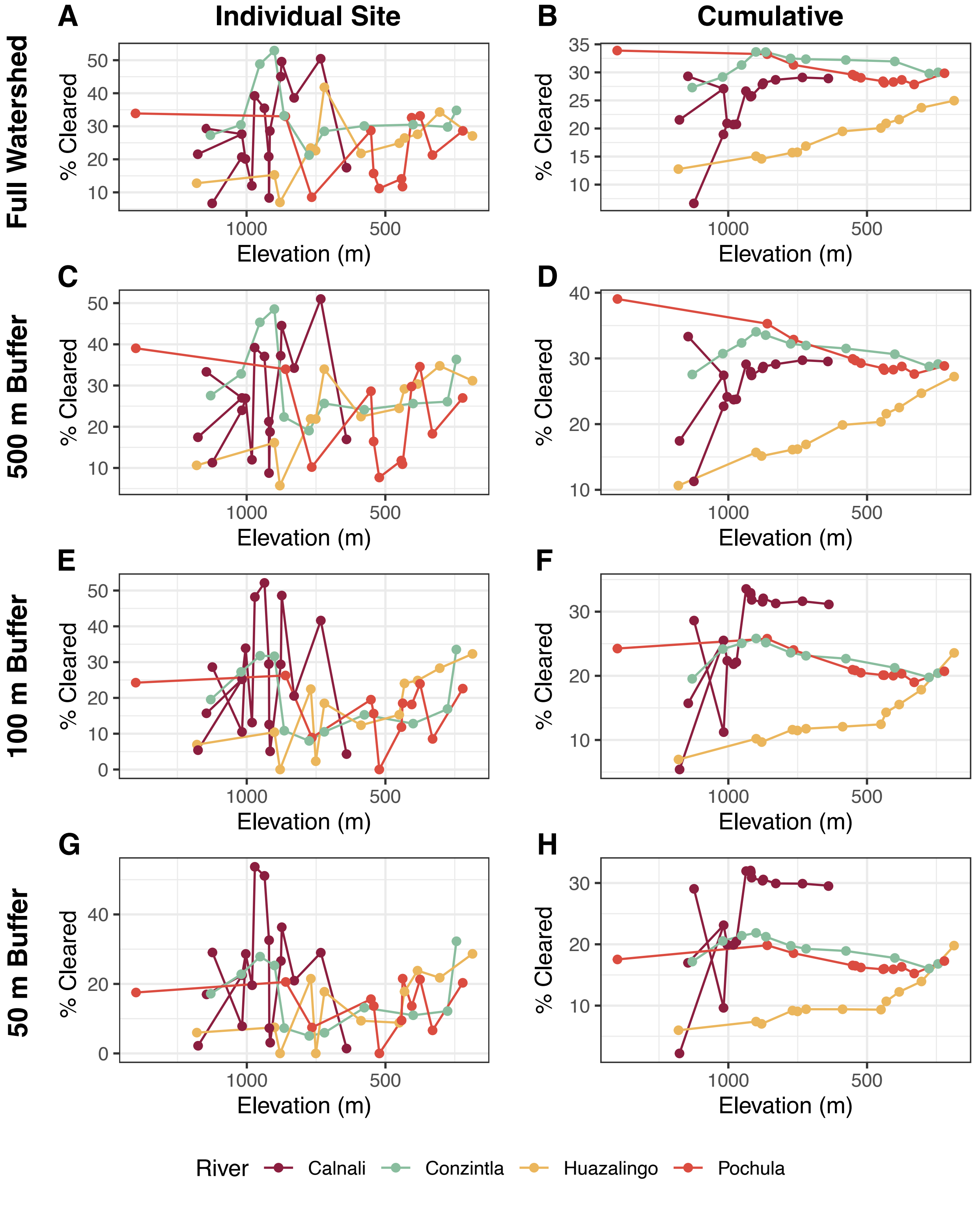


**Figure S2. Cleared land proportions from GIS analysis. Related to Figure 5.**
Proportion of land cover identified as cleared land (non-native vegetation, grasslands, and pasture) in each watershed in 2020–2023 based on analysis in ArcGIS Pro. The left column shows the proportion at each site, based on analysis of the subcatchment

between that site and the site directly upstream (see Figure S4A). The right column shows the cumulative subcatchment of all sections of the stream upstream of a given site. The first row shows calculations incorporating pixels at any distance from a stream within the focal subcatchment area, while lower rows show calculations limited to pixels within **C–D)** 500 meters, **E–F)** 100 meters, or **G–H)** 50 meters of a stream. Streams were designated as pixels with accumulated flow > 3000 cells, depicted with blue lines in Figure 5A. Colors correspond to stream (see legend at bottom), points represent sites, lines connect sites in the order in which they occur on the stream (note that Calnali line branches due to inclusion of multiple headwater tributaries).

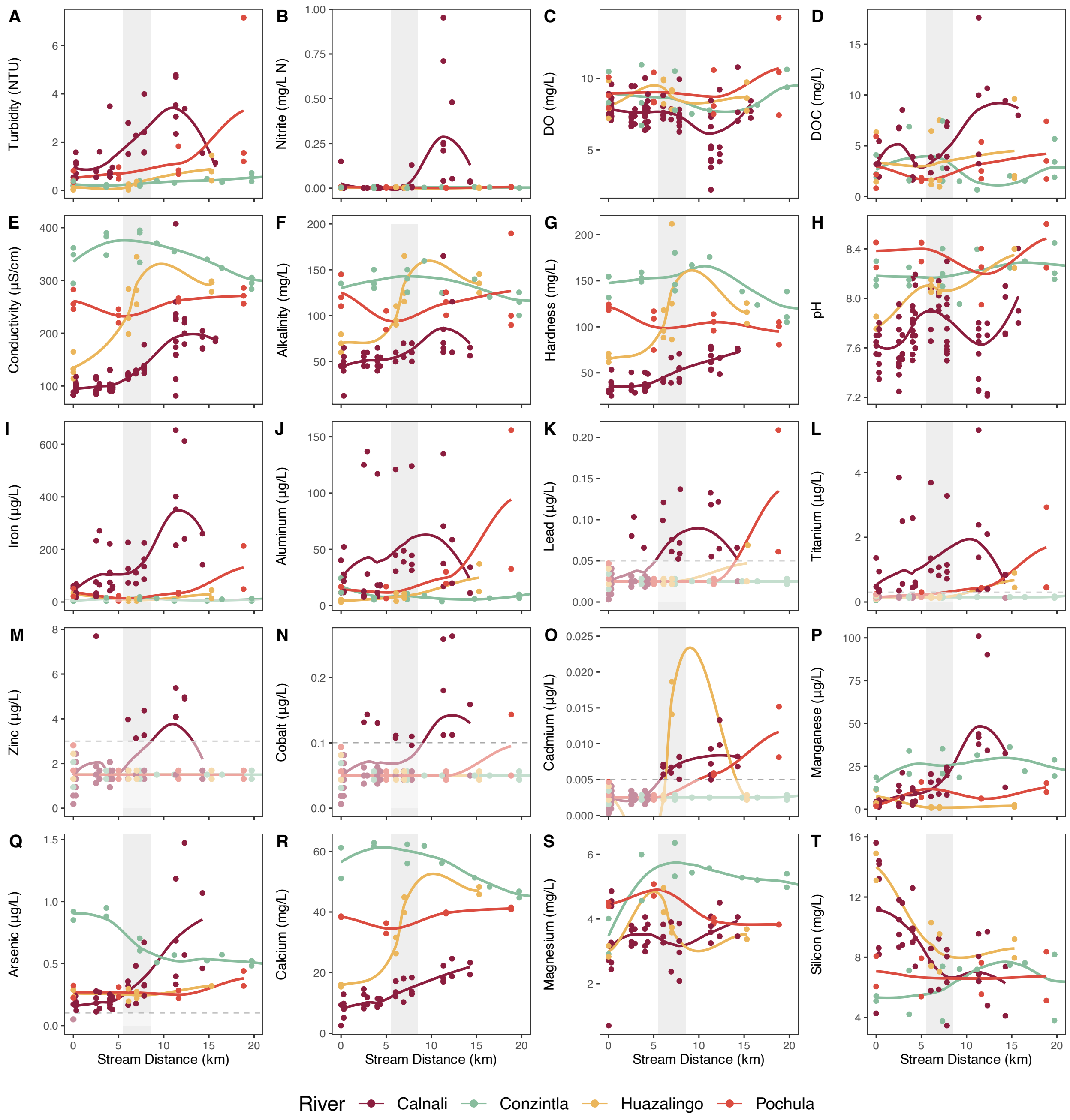


**Figure S3. Plots of water quality parameters along each stream. Related to Figure 6**. Points represent individual samples. Numbers on the X-axis correspond to the position along the stream, with 0 indicating the most upstream site sampled. Values on the Y-axis correspond to measurements of the chemical parameter, and color denotes stream of sampling (see legend at bottom). Lines are LOESS curves fit through data for each stream, and gray bars indicate the span in which the Río Calnali passes through the town of Calnali. Panels represent **A)** turbidity, in nephelometric turbidity units **B)** nitrite, in mg N/L **C)** dissolved oxygen, in mg $O_2$/L **D)** dissolved organic carbon from Shimadzu TOC-L analyzer, in mg C/L **E)** total (special) conductivity, in µS/cm **F)** alkalinity, in mg/L $CaCO_3$ equivalents **G)** total hardness, in mg/L $CaCO_3$ equivalents **H)**

pH **I–T)** ICP-MS measurements, including **I)** Iron **J)** Aluminum **K)** Lead **L)** Titanium **M)** Zinc **N)** Cobalt, **O)** Cadmium **P)** Manganese **Q)** Arsenic **R)** Calcium **S)** Magnesium and **T)** Silicon. Grey shaded area denotes the sites located within the town of Calnali on the Río Calnali transect. Dashed horizontal lines and translucent zones in **K–O)** and **Q)** represent measurements that fell below the detection limit of the ICP-MS assay. Points below this limit are arbitrarily plotted at half the detection threshold, with jittering for visualization. Note that these values below the line do not represent point estimates of the ion concentrations.

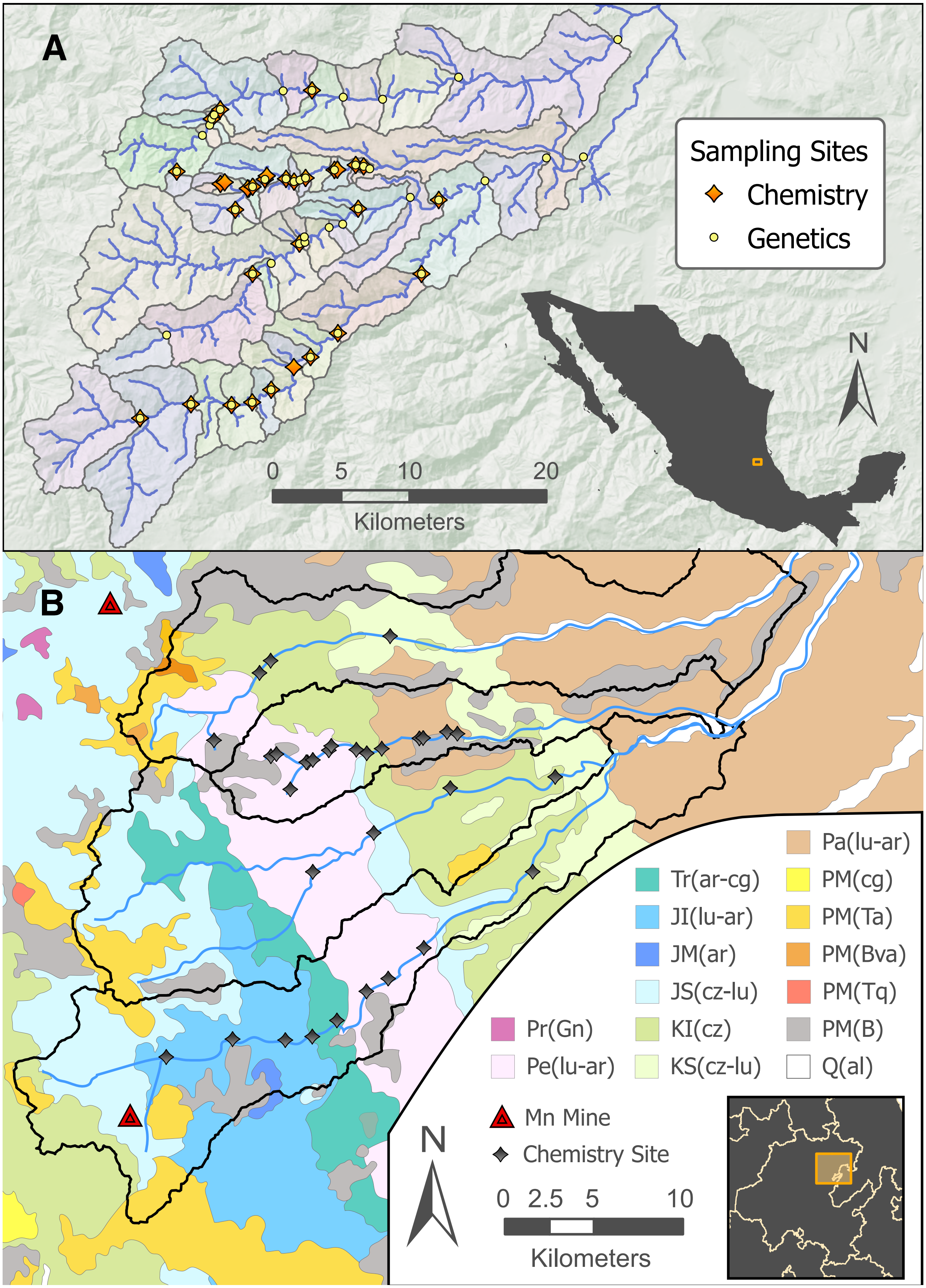
A
Sampling Sites
Chemistry
Genetics
N
0 5 10 20
Kilometers
B
Pa(lu-ar)
Tr(ar-cg)
PM(cg)
JI(lu-ar)
PM(Ta)
JM(ar)
PM(Bva)
JS(cz-lu)
PM(Tq)
Pr(Gn)
KI(cz)
PM(B)
Pe(lu-ar)
KS(cz-lu)
Q(al)
Mn Mine
Chemistry Site
N
0 2.5 5 10
Kilometers

**Figure S4. Hydrology and geology of the Río Atlapexco drainage. Related to Figures 5–6. A)** Locations of sampling sites overlaid with the inferred subcatchment of each site. Streams are shown as blue lines, sites where fish were collected are yellow circles, and sites where water chemistry data were collected are orange diamonds. The subcatchment draining to each site is indicated with arbitrarily colored translucent areas separated by dark lines. Note that the full drainage of each site includes all subcatchments upstream of it (i.e. not only the closest subcatchment, but all the subcatchments of all upstream sites). **B)** Geological features underlying the sampled stream drainages in the Atlapexco Basin. Black outlines denote drainage area for the four different streams included in this study. From top to bottom these streams are the Río Huazalingo, Río Calnali, Río Pochula, and Río Conzintla. Blue lines show streams, gray points show the sites of water chemistry sampling, and red triangles denotes locations of open-pit manganese mines. This geological map was adapted from the Mexican National Institute of Statistics and Geography (INEGI)[S1]. The color represents age and type of bedrock. The abbreviations in the legend include text outside parentheses specifying geological age (Pr – Precambrian; Pe – Permian; Tr – Triassic; JI – Inferior Jurassic; JM – Middle Jurassic; JS – Superior Jurassic; KI – Inferior Cretaceous; KS – Superior Cretaceous; Pa – Paleogene; PM – Pliocene/Miocene; Q – Quaternary) and inside parentheses indicating rock type (al – alluvial soil; ar – sandstone; B – basalt; Bva – acidic volcanic breccia; cg – conglomerate; cz – limestone; Gn – gneiss; lu – mudstone; Ta – acidic tuff; Tq – trachyte). The role of bedrock weathering in determining water chemistry is supported by the loading of major ionic components of bedrock (Si, Ca, Mg, S) onto the same vector in PC space which separates watersheds (Figure 6). The effect of geological variation could also be compounded by the presence of an open-pit manganese mine in the headwaters of the Río Conzintla, which may increase sediment inputs to the stream.

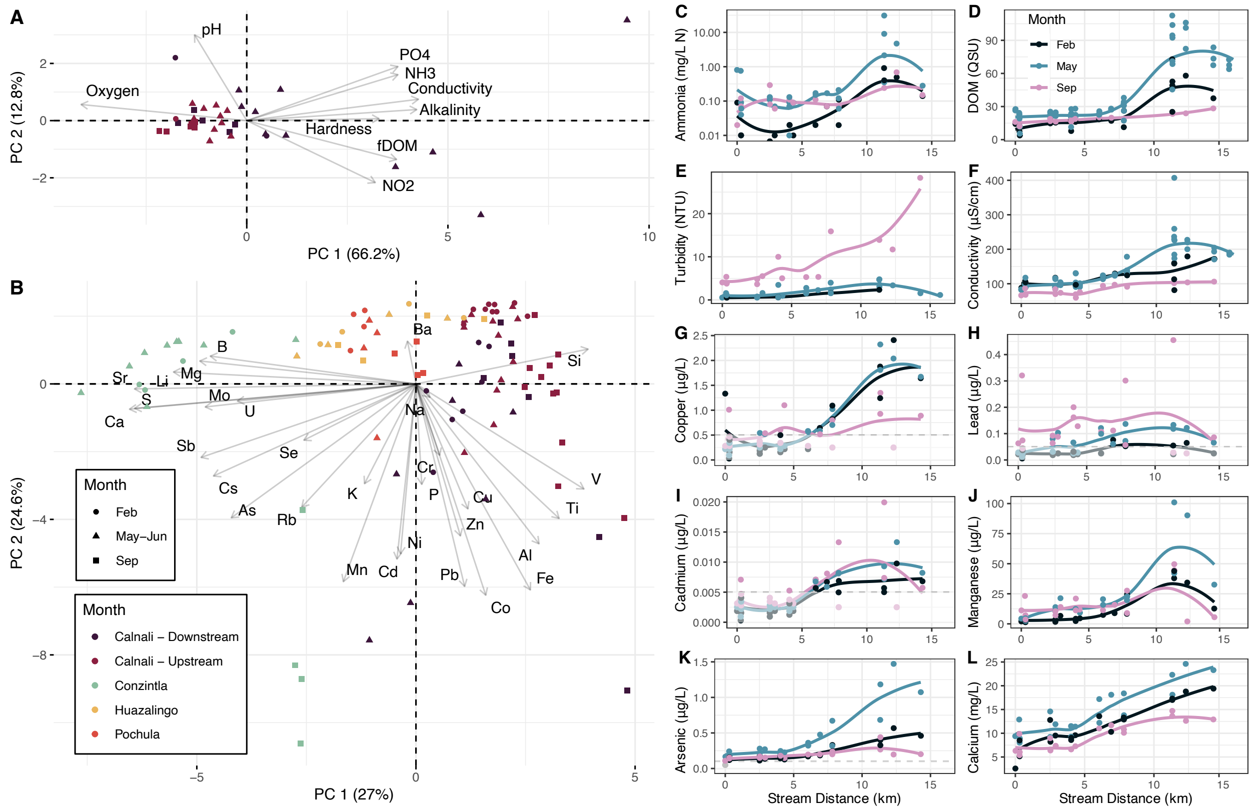

**Figure S5. Water chemistry including September samples. Related to Figure 6.** Principal component analysis of water chemistry data highlighting **A)** temporal variation in the Río Calnali in non-metals data and **B)** ICP-MS metals data from all sites and dates. PCAs were constructed as in main text, except for inclusion of data from the rainy season in September; sulfate measurements were not collected at that timepoint, and thus this parameter was omitted from the non-metals PCA. Points represent individual sampling events, with shape and color denoting sampling month and drainage, respectively. Arrows and text show loading of variables on first two principal components. Proportion of variance explained is indicated parenthetically on the x and y axes. Concentrations of nearly all non-metal parameters were lower in September, and differences between sites upstream and downstream of the town of Calnali were minimized. This suggests that any increase in surface inputs during the rainy season is offset by increased flow, ultimately causing dilution of pollutants. Notably, Conzintla samples from September were collected after an exceptionally large rainfall caused elevated water levels and turbidity. Divergent PC2 from this sampling event suggest this drainage may be especially sensitive to short-term water chemistry fluctuations driven by erosion, potentially due to the open-pit manganese mine present in its headwaters (Figure S4B). **C-L)** Temporal variation in water chemistry along the Río Calnali for various water chemistry parameters. Points represent sampling events, and X-axis indicates sampling site position along the length of the Río Calnali. Zero corresponds to the highest elevation site sampled. Values on the Y-axis correspond to measurements of the chemical parameter sampled, and the color denotes month of sampling. Lines are LOESS curves fit to this data. Dashed line and translucent zones in **G)**, **H)**, **I)**, and **K)**

indicate measured values below the detection limit of the ICP-MS assay. Points below this limit are plotted at half the detection threshold, with jittering to prevent overlap and visually represent sample size, but do not represent point estimates of the concentration. Abbreviations: DOM = fluorescent Dissolved Organic Matter; QSU = quinine sulfate units; NTU = nephelometric turbidity units.

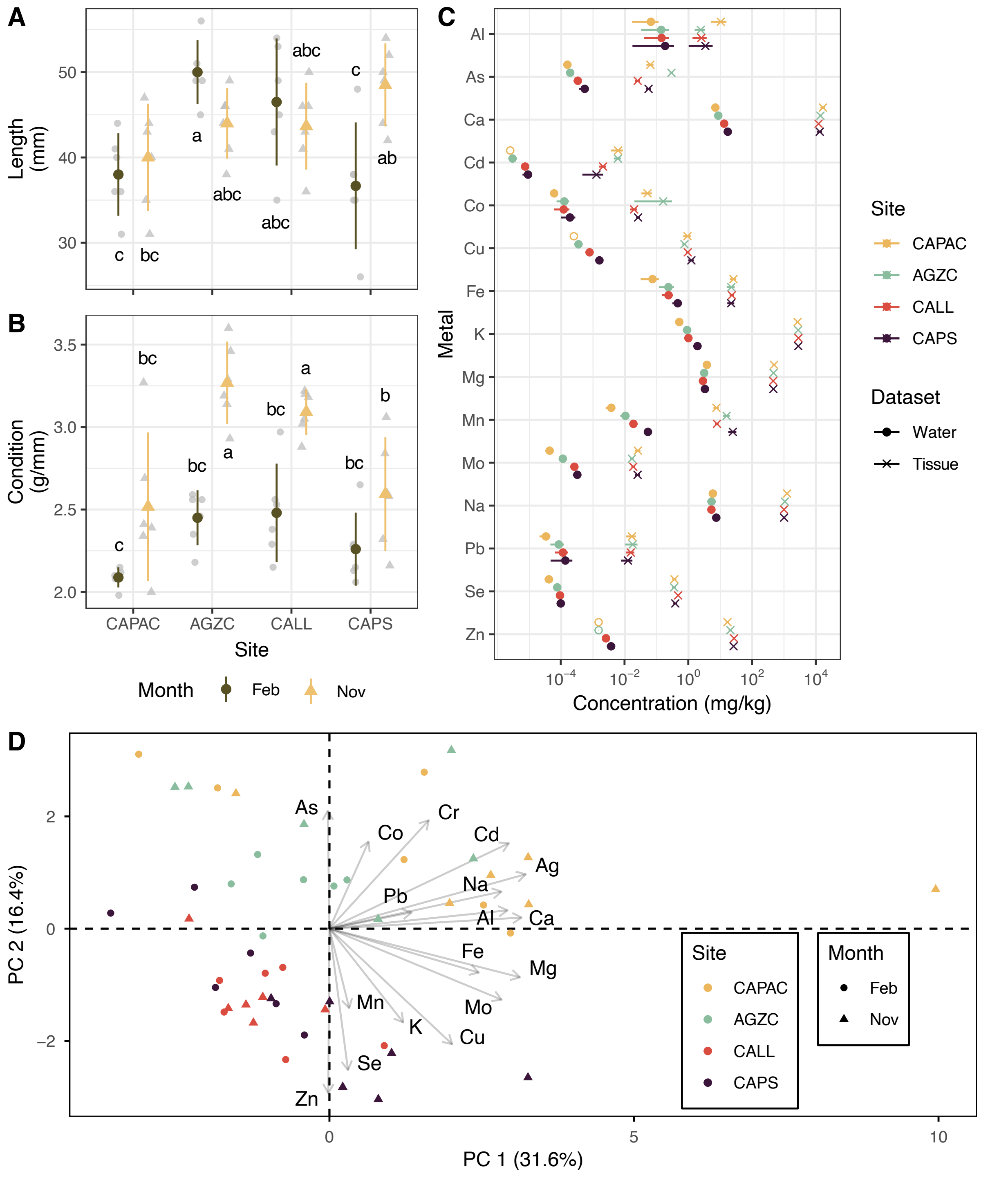


**Figure S6. ICP-MS metal analysis of fish tissue from four sites on the Río Calnali. Related to Figure 6.** Two sites (CAPAC and AGZC) are located upstream of the town of Calnali, and the others (CALL and CAPS) are located downstream of the town. **A–B)** Length (**A**) and condition factor (**B**) of fish collected for ICP-MS, separated by site and collection month. Gray points represent individual samples, colored points and bars represent group mean ± 2 standard errors. Letters denote statistical differences from Tukey's HSD test. Fish length differed by site (ANOVA $P = 0.005$) and its interaction with month ($P = 0.001$), but not month alone ($P = 0.425$) and individual condition varied with site (ANOVA $P = 3.12e\text{-}6$) and month ($P = 4.68e\text{-}9$), but not their interaction ($P = 0.114$). **C)** Concentrations of metals from ICP-MS measurements, compared between water (•) and fish tissue (×). Y-axis separates each metal tested, color denotes sampling site, and points and error bars represent mean ± 2 SE concentration for each site. Error bars are invisible in some cases due to small variance relative to the X-axis scale. Open circles denote sites and metals for which all values were below the detection limit, in which case the point was arbitrarily placed at one half the detection limit. All tissue measurements were above detection limits. **D)** Principal component analysis of metal concentrations from whole fish tissue. Points represent individual sampled fish, with shape and color denoting sampling month and site, respectively. Arrows and text show loading of variables on first two principal components. Proportion of variance explained is indicated parenthetically on the X- and Y-axes. Our results suggest that any mechanistic connection between aqueous metals and reproductive isolation is unlikely to be mediated by gross bioaccumulation of toxic ions in the body.

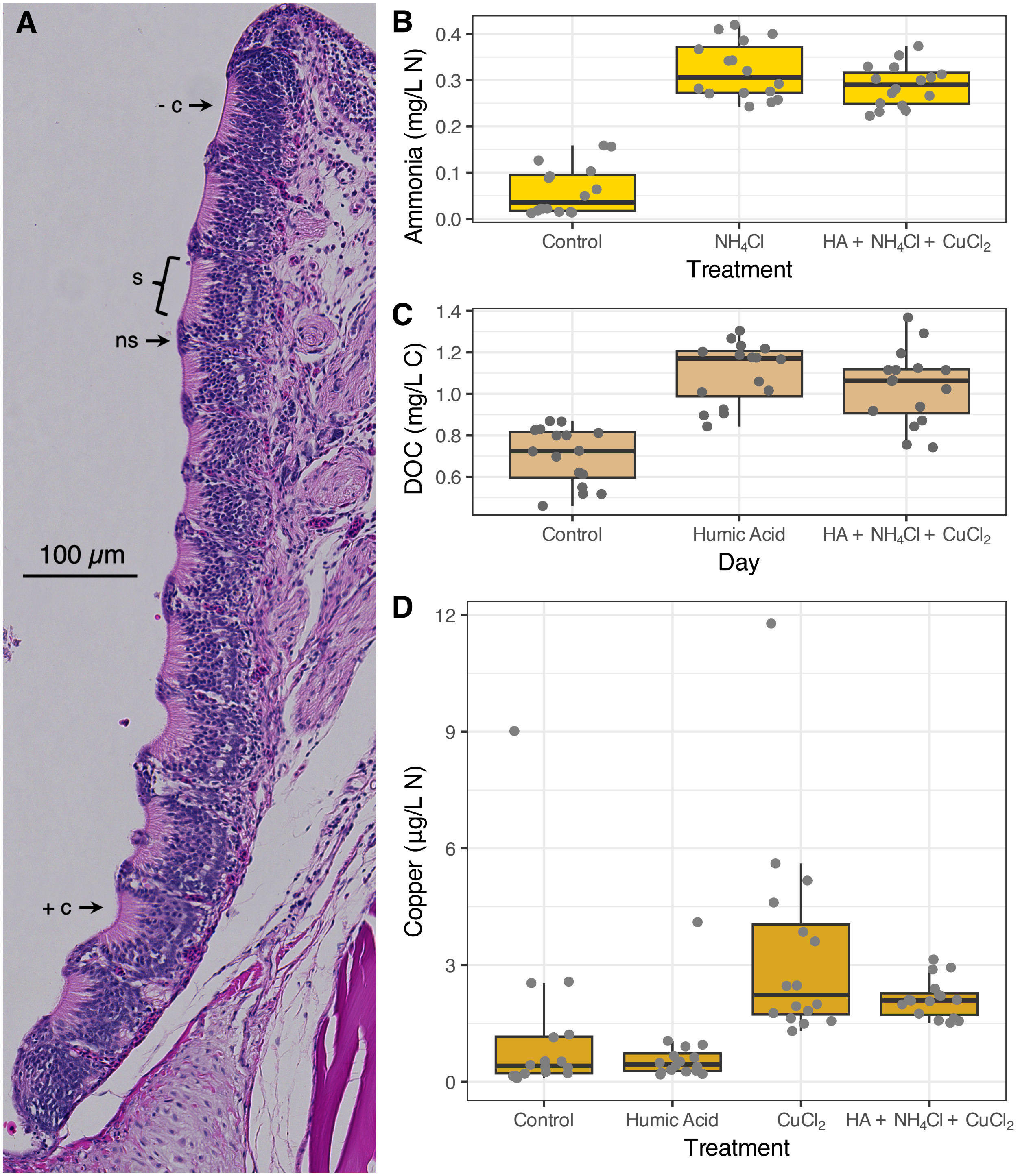


**Figure S7. Olfactory histological example and experimental quality control. Related to Figure 7. A)** Representative image of hematoxylin & eosin (H&E) stained olfactory rosette of *Xiphophorus birchmanni* × *malinche* hybrid. Sensory epithelial beds (s) are visible due to eosinophilic (pink) apical surface of sensory neurons, with basophilic (purple) surface cells visible in the intervening non-sensory epithelial surface (ns). Examples of sensory beds where cilia are absent (- c) or present (+ c) are highlighted. Note that images were taken at 400X magnification with a Cytation 5 slide scanner, but cilia were quantified at 1,000X magnification using a compound microscope. **B–D)** Water chemistry parameters measured during laboratory treatments testing chemical effects on olfactory histology. Individual water samples, which are shown as gray points, were taken from each tank at the start of each trial, before and after daily water changes, and at the end of the trial. **B)** Ammonia, measured as $NH_4$. **C)** Dissolved organic carbon, measured as non-purgeable organic carbon. **D)** Copper measured by ICP-MS. In general, testing was performed on the no-manipulation control treatment and the two treatments in which the tested parameter was directly manipulated. However, the Humic Acid (HA) treatment was also tested for its effect on copper levels due to previous reports of heavy metals in commercially purchased humic acid[S2].

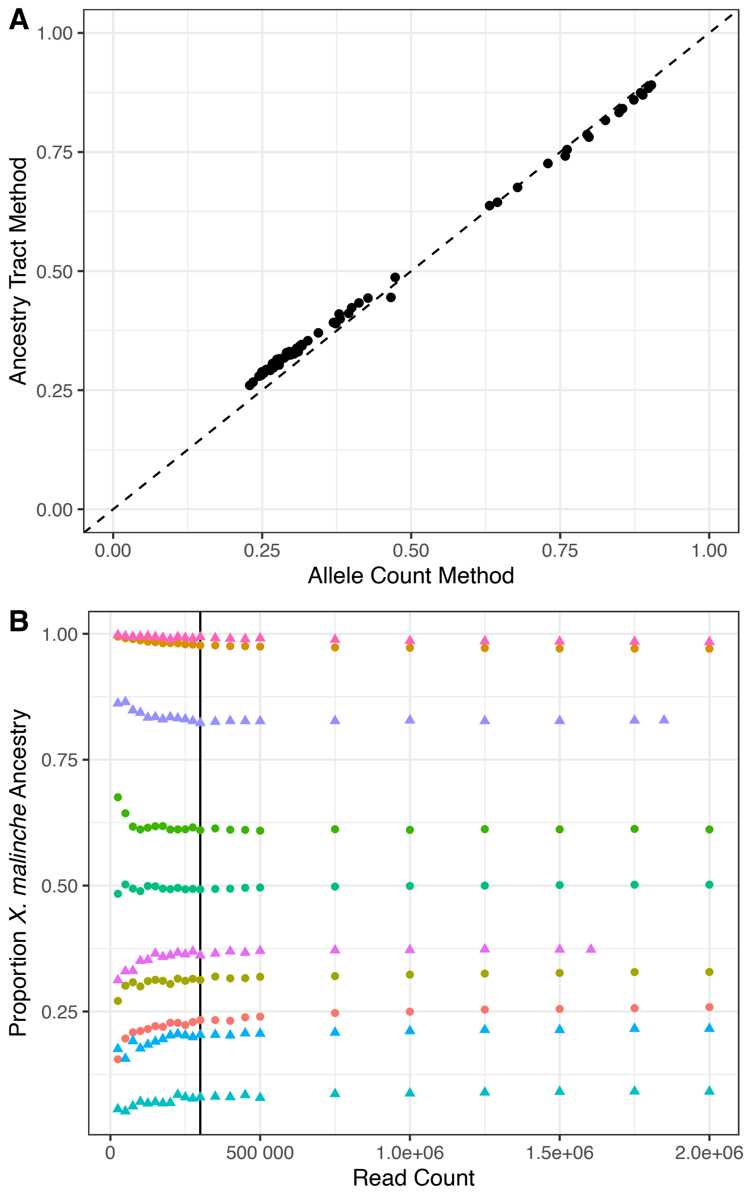


**Figure S8. Ancestry inference methods quality control. Related to Figure 3. A)** Comparison of different approaches for estimating ancestry fraction. Individuals plotted here represent 73 samples from the Plaza (PLAZ) site. On the X-axis, the proportion of the genome derived from *X. malinche* was estimated based on allele counts at ancestry informative markers. On the Y-axis, the proportion of the genome derived from *X. malinche* was estimated using the lengths of ancestry tracts inferred to belong to a particular ancestry state (homozygous *X. malinche*, heterozygous, or homozygous *X. birchmanni*). The dotted line represents the expected 1:1 relationship between these estimation methods. **B)** Downsampling analysis testing sensitivity of genome-wide ancestry fraction estimates to read counts and library type. Five individuals from 150bp paired-end sequencing runs (circles) and five individuals from 100bp single-read runs (triangles) were downsampled to varying coverage before ancestry inference, with the number of reads sampled represented on the X-axis, and the inferred genome-wide ancestry fraction on the Y-axis. Colors distinguish ancestry runs from reads of the same individual, and the vertical line denotes the read count cutoff used in the main text (300,000 reads).

| # | Drainage | Site Name | Site Code | Lat | Lon | Elevation | Genetic | Water |
|---|---|---|---|---|---|---|---|---|
| 1 | Huazalingo | Veneno | VNNO | 20.90358 | -98.6601 | 1180 | T | T |
| 2 | Huazalingo | Cansada | CNSD | 20.926981 | -98.643206 | 900 | T | F |
| 3 | Huazalingo | Cascada | CSCD | 20.933614 | -98.638419 | 880 | T | F |
| 4 | Huazalingo | Corazón | CRZN | 20.93761 | -98.63713 | 769 | T | T |
| 5 | Huazalingo | Totonicapa to Cascade Pool 1 | TCP1 | 20.940231 | -98.635508 | 751 | T | F |
| 6 | Huazalingo | Totonicapa | TOTO | 20.94379 | -98.63151 | 720 | T | T |
| 7 | Huazalingo | Acahuasco Low | AHLW | 20.955858 | -98.59015 | 588 | T | F |
| 8 | Huazalingo | Acuapa | ACUA | 20.95618 | -98.57121 | 450 | T | T |
| 9 | Huazalingo | Pilcuatla | PILC | 20.951744 | -98.550669 | 431 | T | F |
| 10 | Huazalingo | San Pedro | SNPD | 20.950418 | -98.524769 | 384 | T | F |
| 11 | Huazalingo | Tlatzonco | TLZO | 20.964663 | -98.474852 | 304 | T | F |
| 12 | Huazalingo | Achiquihuixtla | ACHI | 20.989106 | -98.370194 | 186 | T | F |
| 13 | Calnali | Calnali High South | CAHS | 20.89553 | -98.63139 | 1176 | F | T |
| 14 | Calnali | Calnali High | CALH | 20.8966 | -98.62859 | 1145 | F | T |
| 15 | Calnali | CAPASC Fuente de Agua | CAPAC | 20.87878 | -98.62159 | 1124 | T | T |
| 16 | Calnali | Calnali Branch South | CABS | 20.8924 | -98.61285 | 1017 | F | T |
| 17 | Calnali | Calnali Branch North | CABN | 20.89252 | -98.61352 | 1017 | F | T |
| 18 | Calnali | Calnali Mid | CALM | 20.89366 | -98.61024 | 1004 | T | T |
| 19 | Calnali | Aguazarca | AGZC | 20.8985 | -98.60215 | 981 | T | T |
| 20 | Calnali | Plank | PLNK | 20.90074 | -98.60091 | 971 | F | T |
| 21 | Calnali | Piloncillo | PILO | 20.89883 | -98.58828 | 936 | T | T |
| 22 | Calnali | Plaza | PLAZ | 20.89704 | -98.58302 | 920 | T | T |
| 23 | Calnali | Peatonal | PEAT | 20.898047 | -98.579144 | 919 | T | F |
| 24 | Calnali | Calnali Low | CALL | 20.89936 | -98.57544 | 916 | T | T |
| 25 | Calnali | Tlalica | CAPS | 20.90459 | -98.55644 | 877 | T | T |
| 26 | Calnali | Tlalica Down | TLCD | 20.90475 | -98.55456 | 874 | F | T |
| 27 | Calnali | Pezmatlán | PEZM | 20.90778 | -98.5425 | 829 | T | T |
| 28 | Calnali | Chahuaco Falls | CHAF | 20.90689 | -98.53726 | 733 | T | T |
| 29 | Calnali | Gilroy | GILR | 20.905331 | -98.533239 | 640 | F | T |
| 30 | Pochula | Culhuacán | CULH | 20.797511 | -98.666703 | 1400 | T | F |
| 31 | Pochula | Texcaco | TEXC | 20.83716 | -98.61026 | 860 | T | T |
| 32 | Pochula | Cascada Real | CSRL | 20.844025 | -98.597997 | 765 | T | F |
| 33 | Pochula | Pochula | POCH | 20.85677 | -98.57946 | 552 | T | T |
| 34 | Pochula | Pochula Low | PHLW | 20.857669 | -98.575858 | 542 | T | F |
| 35 | Pochula | Ocotepec | OCOT | 20.861167 | -98.576208 | 522 | T | F |
| 36 | Pochula | Islet | ISLT | 20.867308 | -98.559903 | 442 | T | F |
| 37 | Pochula | Pico de Aguila | PICO | 20.869431 | -98.550997 | 438 | T | F |

| | | | | | | | | |
|---|---|---|---|---|---|---|---|---|
| *38* | Pochula | Tula | TULA | 20.87946 | -98.54086 | 405 | T | T |
| *39* | Pochula | Bicho | VCHO | 20.886817 | -98.506633 | 375 | T | F |
| *40* | Pochula | Atempa | ATEM | 20.8851 | -98.4879 | 330 | T | T |
| *41* | Pochula | Papatlatla | PAPA | 20.8975 | -98.4575 | 276 | T | F |
| *42* | Conzintla | Coatitlamixtla | CTMX | 20.74362 | -98.6842 | 1130 | T | T |
| *43* | Conzintla | Xochicoatlán | XOCH | 20.75265 | -98.65074 | 1020 | T | T |
| *44* | Conzintla | Mixtla | MXLA | 20.75217 | -98.62412 | 952 | T | T |
| *45* | Conzintla | Mazahuacán | MZHC | 20.75393 | -98.6105 | 900 | T | T |
| *46* | Conzintla | Tlaxcoya | TLCY | 20.76208 | -98.59806 | 865 | T | T |
| *47* | Conzintla | La Esperanza | LESP | 20.77676 | -98.58318 | 775 | F | T |
| *48* | Conzintla | Papaxtla | PPXL | 20.78322 | -98.5722 | 720 | T | T |
| *49* | Conzintla | Otlamalacatla | OMCA | 20.79871 | -98.55421 | 576 | T | T |
| *50* | Conzintla | Zacatipán | ZACA | 20.837186 | -98.499236 | 400 | T | T |
| *51* | Pochula | Huiznopala | HZNP | 20.912376 | -98.417092 | 244 | T | F |
| *52* | Pochula | El Arenal | AREN | 20.913033 | -98.392794 | 221 | T | F |
| *53* | Cal Trib | Calnali Tributary Up | CATU | 20.902586 | -98.573572 | 926 | T | F |
| *54* | Cal Trib | Calnali Tributary Middle | CATM | 20.904600 | 98.569500 | 925 | T | F |
| *55* | Cal Trib | Calnali Tributary Down | CATD | 20.907700 | 98.565900 | 929 | T | F |
| *56* | Cal Trib | Ciudad de Ranas | CDDR | 20.905225 | -98.557397 | 892 | T | F |

**Table S1. List of sampling locations included in this study. Related to Figure 1.** Lat – Latitude, Lon – Longitude. T indicates that genetic data or water chemistry data was collected for that site, F indicates that it was not.

| Drainage | Site Code | Year | # Collected |
|---|---|---|---|
| *Huazalingo* | VNNO | 2014 | 40 |
| | CNSD | 2015 | 26 |
| | CSCD | 2014 | 19 |
| | CRZN | 2014 | 28 |
| | | 2021 | 46 |
| | | 2024 | 39 |
| | TCP1 | 2014 | 31 |
| | TOTO | 2013 | 16 |
| | | 2014 | 120 |
| | | 2015 | 91 |
| | AHLW | 2016 | 19 |
| | ACUA | 2013 | 36 |
| | | 2015 | 13 |
| | PILC | 2013 | 32 |
| | SNPD | 2013 | 22 |
| | TLZO | 2013 | 28 |
| | | 2014 | 10 |
| | ACHI | 2014 | 15 |
| | | 2016 | 4 |
| *Calnali* | CAPAC | 2017 | 23 |
| | CALM | 2021 | 79 |
| | AGZC | 2016 | 51 |
| | PILO | 2012 | 26 |
| | | 2017 | 54 |
| | PLAZ | 2017 | 73 |
| | PEAT | 2017 | 37 |
| | CALL | 2017 | 234 |
| | | 2018 | 74 |
| | | 2020 | 38 |
| | | 2021 | 201 |
| | | 2022 | 155 |
| | CAPS | 2021 | 12 |
| | | 2022 | 24 |
| | PEZM | 2016 | 22 |
| | CHAF | 2017 | 50 |
| | | 2018 | 138 |
| | | 2019 | 60 |
| | GILR | 2019 | 23 |

| Drainage | Site Code | Year | # Collected |
|---|---|---|---|
| *Pochula* | CULH | 2015 | 26 |
| | TEXC | 2015 | 23 |
| | CSRL | 2015 | 16 |
| | POCH | 2015 | 17 |
| | | 2016 | 22 |
| | PHLW | 2016 | 19 |
| | OCOT | 2016 | 26 |
| | ISLT | 2016 | 36 |
| | PICO | 2016 | 39 |
| | TULA | 2015 | 33 |
| | VCHO | 2016 | 14 |
| | | 2017 | 31 |
| | ATEM | 2015 | 14 |
| | | 2017 | 23 |
| | PAPA | 2006 | 4 |
| | | 2008 | 9 |
| *Conzintla* | CTMX | 2023 | 54 |
| | XOCH | 2018 | 54 |
| | | 2021 | 16 |
| | MXLA | 2022 | 29 |
| | MZHC | 2023 | 17 |
| | TLCY | 2023 | 12 |
| | PPXL | 2023 | 19 |
| | OMCA | 2023 | 19 |
| | ZACA | 2019 | 19 |
| | HZNP | 2006 | 24 |
| | | 2007 | 2 |
| | AREN | 2016 | 39 |
| *Calnali* | CATU | 2020 | 6 |
| *Tributary* | CATM | 2020 | 25 |
| | CATD | 2020 | 31 |
| | CDDR | 2021 | 12 |

**Table S2. List of genetic sample sizes by location and date. Related to Figure 3.** Site codes as in Table S1.

| *Drainage* | *Site Code* | *Sample Size* | *Mean Ancestry* | *Std. Dev. Ancestry* | D *Statistic* | P*-value* | *Bonferroni Significance* |
|---|---|---|---|---|---|---|---|
| *Calnali* | CAPAC | 23 | 0.982 | 0.002 | 0.0632119 | 0.664125 | |
| | CALM | 79 | 0.647 | 0.347 | 0.1597263 | 0 | * |
| | AGZC | 51 | 0.584 | 0.351 | 0.2146567 | 0 | * |
| | PILO | 80 | 0.490 | 0.284 | 0.1161647 | 0 | * |
| | PLAZ | 73 | 0.414 | 0.215 | 0.0453731 | 0.320643 | |
| | PEAT | 37 | 0.391 | 0.184 | 0.0637864 | 0.274838 | |
| | CALL | 702 | 0.448 | 0.190 | 0.0108287 | 0.896476 | |
| | CAPS | 36 | 0.503 | 0.148 | 0.0692381 | 0.182777 | |
| | PEZM | 22 | 0.628 | 0.154 | 0.0617895 | 0.737224 | |
| | CHAF | 248 | 0.686 | 0.105 | 0.0146906 | 0.990367 | |
| | GILR | 23 | 0.689 | 0.082 | 0.0637874 | 0.647471 | |
| *Conzintla* | CTMX | 54 | 0.996 | 0.010 | 0.0328751 | 0.965143 | |
| | XOCH | 70 | 0.594 | 0.438 | 0.1924042 | 0 | * |
| | MXLA | 29 | 0.374 | 0.383 | 0.1281781 | 0.000211 | * |
| | MZHC | 17 | 0.090 | 0.018 | 0.0773580 | 0.538903 | |
| | TLCY | 12 | 0.117 | 0.084 | 0.0870907 | 0.599963 | |
| | PPXL | 19 | 0.082 | 0.016 | 0.0567116 | 0.916779 | |
| | OMCA | 19 | 0.085 | 0.013 | 0.0502825 | 0.980973 | |
| | ZACA | 19 | 0.169 | 0.015 | 0.0800843 | 0.385644 | |
| | HZNP | 26 | 0.162 | 0.017 | 0.0569652 | 0.747239 | |
| | AREN | 39 | 0.174 | 0.013 | 0.0490792 | 0.693023 | |
| *Huazalingo* | VNNO | 40 | 0.986 | 0.002 | 0.0405727 | 0.919785 | |
| | CNSD | 26 | 0.984 | 0.002 | 0.0540566 | 0.821784 | |
| | CSCD | 19 | 0.988 | 0.014 | 0.1129799 | 0.032045 | |
| | CRZN | 113 | 0.343 | 0.213 | 0.0319384 | 0.586268 | |
| | TCP1 | 31 | 0.246 | 0.116 | 0.0512672 | 0.78897 | |
| | TOTO | 227 | 0.245 | 0.117 | 0.0161153 | 0.98159 | |
| | AHLW | 19 | 0.208 | 0.021 | 0.0468443 | 0.991205 | |
| | ACUA | 49 | 0.202 | 0.025 | 0.0373504 | 0.91452 | |
| | PILC | 32 | 0.101 | 0.016 | 0.0491217 | 0.825054 | |
| | SNPD | 22 | 0.088 | 0.022 | 0.0793971 | 0.29282 | |
| | TLZO | 38 | 0.081 | 0.008 | 0.0501598 | 0.675376 | |
| | ACHI | 19 | 0.074 | 0.007 | 0.1045109 | 0.067967 | |
| *Pochula* | CULH | 26 | 0.998 | 0.001 | 0.0603902 | 0.648209 | |
| | TEXC | 23 | 0.999 | 0.000 | 0.0608561 | 0.72945 | |
| | CSRL | 16 | 0.998 | 0.000 | 0.0614887 | 0.914394 | |
| | POCH | 39 | 0.994 | 0.009 | 0.0370263 | 0.977026 | |
| | PHLW | 19 | 0.995 | 0.005 | 0.0516198 | 0.970474 | |

| | | | | | |
|---|---|---|---|---|---|
| OCOT | 26 | 0.948 | 0.060 | 0.0572164 | 0.740296 |
| ISLT | 36 | 0.635 | 0.119 | 0.0459761 | 0.839922 |
| PICO | 39 | 0.520 | 0.076 | 0.0307243 | 0.993883 |
| TULA | 33 | 0.474 | 0.019 | 0.0427192 | 0.946834 |
| VCHO | 45 | 0.420 | 0.027 | 0.0380131 | 0.929538 |
| ATEM | 37 | 0.337 | 0.047 | 0.0509313 | 0.66973 |
| PAPA | 13 | 0.211 | 0.020 | 0.0584610 | 0.984087 |

**Table S3. Ancestry statistics for genetic sampling sites. Related to Figure 3.** Mean and standard deviation in ancestry are based on individual measurements of the proportion of ancestry informative markers derived from *X. malinche*, with 0 indicating pure *X. birchmanni* and 1 indicating pure *X. malinche*. *D* statistic and *P*-value are from Hartigan's Dip Test for unimodality. Asterisks denote sites that significantly departed from unimodality after Bonferroni correction for multiple testing. Sites are ordered upstream to downstream within each drainage.

| Drainage | Site Code | Full Sample N | Full Sample D | Mean Down-sampled D | Proportion $P < 0.05$ | Proportion Bonferroni Significant |
|---|---|---|---|---|---|---|
| *Huazalingo* | CRZN | 113 | 0.0319384 | 0.061501 | 0.014 | 0 |
| | TOTO | 227 | 0.0161153 | 0.058308 | 0 | 0 |
| *Calnali* | CALM | 79 | 0.1597263 | 0.162670 | 0.967 | 0.845 |
| | AGZC | 51 | 0.2146567 | 0.194172 | 0.999 | 0.976 |
| | PILO | 80 | 0.1161647 | 0.125770 | 0.773 | 0.429 |
| | PLAZ | 73 | 0.0453731 | 0.068764 | 0.047 | 0.001 |
| | CALL | 702 | 0.0108287 | 0.063760 | 0.02 | 0.001 |
| | CAPS | 36 | 0.0692381 | 0.082225 | 0.095 | 0 |
| | PEZM | 22 | 0.0617895 | 0.061790 | 0 | 0 |
| *Conzintla* | XOCH | 70 | 0.1924042 | 0.186371 | 0.993 | 0.908 |
| | MXLA | 29 | 0.1281781 | 0.128963 | 0.9 | 0.285 |
| *Pochula* | TULA | 33 | 0.0427192 | 0.056138 | 0 | 0 |

**Table S4. Down-sampling analysis of sensitivity to sample size. Related to Figure 3.** Results of down-sampling the number of individuals per population on Hartigan's Dip Statistic results, based on 1,000 replicate simulations. For each simulation, 22 individuals were drawn from the total sample for the population without replacement before calculating Hartigan's Dip Statistic. Rightmost columns represent the proportion of the 1,000 simulations in which the *P*-value for the departure from unimodality was nominally significant (Proportion $P < 0.05$) or significant at the same Bonferroni correction threshold used in the main text (Proportion Bonferroni Significant, i.e. $P < 0.001105$).

**Ammonia**

| ***ANOVA*** | *Sum Sq* | *D.f.* | F | P *(>F)* | *Signif.* |
|---|---|---|---|---|---|
| *Drainage* | 112.784 | 4 | 16.666 | 1.04E-09 | *** |
| *Month* | 122.055 | 2 | 36.072 | 1.40E-11 | *** |
| *Interaction* | 19.024 | 5 | 2.249 | 0.05851 | . |
| *Residuals* | 121.811 | 72 | | | |

| ***Tukey HSD*** | *Group* |
|---|---|
| Huazalingo | A |
| Calnali - Upstream | A |
| Calnali - Downstream | B |
| Pochula | A |
| Conzintla | A |

**Nitrite**

| ***ANOVA*** | *Sum Sq* | *D.f.* | F | P *(>F)* | *Signif.* |
|---|---|---|---|---|---|
| *Drainage* | 100.639 | 4 | 21.867 | 7.67E-12 | *** |
| *Month* | 18.44 | 2 | 8.0135 | 0.000721 | *** |
| *Interaction* | 7.502 | 5 | 1.3041 | 0.271867 | |
| *Residuals* | 82.842 | 72 | | | |

| ***Tukey HSD*** | *Group* |
|---|---|
| Huazalingo | A |
| Calnali - Upstream | A |
| Calnali - Downstream | B |
| Pochula | A |
| Conzintla | A |

**fDOM**

| ***ANOVA*** | *Sum Sq* | *D.f.* | F | P *(>F)* | *Signif.* |
|---|---|---|---|---|---|
| *Drainage* | 45.34 | 4 | 41.695 | <1E-15 | *** |
| *Month* | 6.959 | 2 | 12.800 | 8.91E-06 | *** |
| *Interaction* | 1.054 | 6 | 0.6463 | 0.693 | |
| *Residuals* | 33.438 | 123 | | | |

| ***Tukey HSD*** | *Group* |
|---|---|
| Huazalingo | A |
| Calnali - Upstream | B |
| Calnali - Downstream | B |
| Pochula | A |
| Conzintla | A |

**Turbidity**

| ***ANOVA*** | *Sum Sq* | *D.f.* | F | P *(>F)* | *Signif.* |
|---|---|---|---|---|---|
| *Drainage* | 53.89 | 4 | 23.8405 | 9.16E-12 | *** |
| *Month* | 1.744 | 2 | 1.5433 | 2.22E-01 | |
| *Interaction* | 1.142 | 6 | 0.3367 | 0.9148 | |
| *Residuals* | 33.341 | 59 | | | |

| ***Tukey HSD*** | *Group* |
|---|---|
| Huazalingo | A |
| Calnali - Upstream | B |
| Calnali - Downstream | C |
| Pochula | B |
| Conzintla | A |

**Table S5. Statistical Testing of Water Chemistry. Related to Figure 6.** Results of Type-II ANOVA and Tukey HSD test for differences in water chemistry parameters between streams, with the Río Calnali split into sites upstream and downstream of the center of the town of Calnali. Sites from CAPAC to PILO were treated as upstream, sites from PLAZ to CHAF were treated as downstream. All values were log-transformed to better fit the assumption of normality (Shapiro-Wilks test $P > 0.01$ for all model residuals). Because of the presence of zero values in these measurements, 0.001 was added to ammonia and nitrite values to avoid undefined log-transformations. Drainages with the same letter code were not significantly different by Tukey HSD testing.
